# Repetitive DNA profiles Reveal Evidence of Rapid Genome Evolution and Reflect Species Boundaries in Ground Beetles

**DOI:** 10.1101/2020.01.03.894527

**Authors:** John S. Sproul, Lindsey M. Barton, David R. Maddison

**Affiliations:** Department of Integrative Biology, Oregon State University, 3029 Cordley Hall, Corvallis, Oregon, 97331, USA; Department of Biology, University of Rochester, 402 Hutchison Hall, PO Box 270211, Rochester, New York, 14627, USA

**Keywords:** copy number variation profiles, ribosomal DNA, rapid genome evolution, species delimitation, *Carabidae*, *Bembidion*

## Abstract

Genome architecture is a complex, multidimensional property of an organism defined by the content and spatial organization of the genome’s component parts. Comparative study of entire genome architecture in model organisms is shedding light on mechanisms underlying genome regulation, evolution, and diversification; but such studies require costly analytical approaches which make extensive comparative study impractical for most groups. However, lower-cost methods that measure a single architectural component (e.g., distribution of one class of repeats) have potential as a new data source for evolutionary studies insofar as that measure correlates with more complex biological phenomena, and for which it could serve as part of an explanatory framework. We investigated copy number variation (CNV) profiles in ribosomal DNA (rDNA) as a simple measure reflecting the distribution of rDNA subcomponents across the genome. We find that signatures present in rDNA CNV profiles strongly correlate with species boundaries in the *breve* species group of *Bembidion*, and vary across broader taxonomic sampling in *Bembidion* subgenus *Plataphus*. Profiles of several species show evidence of re-patterning of rDNA-like sequences throughout the genome, revealing evidence of rapid genome evolution (including among sister pairs) not evident from analysis of traditional data sources such as multi-gene data sets. Major re-patterning of rDNA-like sequences has occurred frequently within the evolutionary history of *Plataphus*. We confirm that CNV profiles represent an aspect of genomic architecture (i.e., the linear distribution of rDNA components across the genome) via fluorescence *in-situ* hybridization. In at least one species, novel rDNA-like elements are spread throughout all chromosomes. We discuss the potential of copy number profiles of rDNA, or other repeats, as a low-cost tool for incorporating signal of genomic architecture variation in studies of species delimitation and genome evolution.

## Introduction

Genome architecture is a complex, multidimensional property of an organism (Lynch and Walsh 2007). At the highest levels, genome architecture comprises the spatial organization and content of a genome’s component parts. A genome’s spatial organization encompasses both the relative linear organization within chromosomes of different sequence types, as well as the spatial layout of the genome within the nucleus; the latter is largely driven by DNA-binding protein interactions (Zalensky 1998; Lynch and Walsh 2007; Di Pierro et al. 2017; MacPherson et al. 2018). The component parts of a genome belong to classes of sequences (e.g., coding, intergenic, repetitive, telomeric, centromeric, origins of replication), which themselves have their own regional architecture defined by their subcomponents (e.g., the abundance and organization of specific repeats, exon/intron layout), and their interactions with different classes of DNA-binding proteins and protein complexes. Thus, overall genome architecture arises from a multi-tiered network of DNA-DNA and DNA-protein interactions within the nucleus as constrained by the genome’s linear organization, and the details of that architecture are central to genome stability, DNA repair, gene regulation, DNA replication, and many other processes (Lynch and Walsh 2007).

The re-patterning of architectural components is increasingly identified as a driver of genome evolution and speciation (Kazazian 2004; Feschotte 2008; Biémont 2010; Hall et al. 2016). For example, rapid expansion of specific transposable elements (Stankiewicz and Lupski 2002; Kapusta et al. 2017), expansion and contraction of protein-coding gene families (Koonin 2009), and changes to methylation signatures that affect chromatin structure and gene expression (Madlung et al. 2002; Di Pierro et al. 2017) are all examples of changes within a single component of genome architecture driving genome differentiation and phenotype evolution among lineages.

Research that could benefit from a comparative study of genome architectures can be very costly. Documenting the entire genome architecture of a single specimen is challenging as it entails mapping both the genomic position of sequence classes, and their interactions within and among other classes of sequences and protein classes, a process that requires a combination of costly analytical approaches (e.g., whole-genome sequencing and annotation, HI-C, CHiP Seq, cytogenetic experiments) (Pinkel et al. 1988; Consortium 2002; Krzywinski et al. 2009; Di Pierro et al. 2017); extending this to the multiple specimens and multiple species needed for evolutionary studies can make the research prohibitively expensive.

However, for some research questions in evolutionary genomics, low-cost measures of one component of the genomic architecture might fortuitously provide a signal that captures key aspects of the architecture and offer a powerful lens to understand evolutionary history. The usefulness of any simple, one-dimensional measure of something as complex as genome architecture will depend upon how much that measure correlates with more complex biological phenomena, and for which it could serve as part of an explanatory framework.

In this study, we explore whether the copy number variation (CNV) profile in ribosomal DNA (rDNA), a simple measure reflecting the distribution and abundance of rDNA subcomponents across the genome, is correlated with current and past patterns of gene flow within a suite of species. Our study system is the *Bembidion breve* species group, a small group of closely related ground beetle (Carabidae) species living in montane areas of western North America. In a previous study, we found preliminary evidence that substantial CNV within sequences of rDNA, easily measured through low-coverage genome sequencing, is present across some species in the group (Sproul and Maddison 2017). The copy number (CN) differences between species were sufficiently large as to suggest variation in rDNA repeats could account for genome-scale differences in repeat content between closely related species. For example, in one specimen of *Bembidion laxatum*, 0.6% of all reads obtained through whole-genome shotgun sequencing mapped to the rDNA cistron (the tandemly repeated region of rDNA containing 18S and 28S genes). In contrast, for a specimen of *B*. *lividulum* (a species extremely similar morphologically to *B. laxatum*, Fig. S1), an astounding 16.9% of all genomic reads obtained mapped to the rDNA cistron – the vast majority mapping to a region of the internal transcribed spacers (ITS) and 28S rRNA gene; that region showed dramatic CN inflation relative to other rDNA regions (e.g., the 18S rRNA gene just a few thousand bases upstream). This suggested that patterns of CNV in rDNA could be a simple measure of an aspect of genomic architecture providing insight into genome evolution and speciation, and could be strongly correlated with species boundaries.

Ribosomal DNA occurs in tandem arrays in the highly transcribed nucleolar organizing regions of the genome, with clusters often appearing on more than one chromosome (McClintock 1934; White 1977; Schwarzacher and Wachtler 1993). However, numerous studies document the transfer of rDNA fragments from nucleolar organizing regions into heterochromatin (tightly packed, gene-poor, repeat-rich DNA) where they can undergo extensive multiplication, and subsequent sequence divergence from functional rDNA (McClintock 1934; White 1977; Schwarzacher and Wachtler 1993; Martins et al. 2006; Raskina et al. 2008; Nguyen et al. 2010; Cioffi and Bertollo 2012; Iwata-Otsubo et al. 2016). These mobilized fragments of rDNA can be thought of newly birthed, rDNA-like repetitive elements. Mobilization of rDNA has been documented using cytogenetic methods in many groups including plants (Raskina et al. 2004; Qi et al. 2015; Ding et al. 2016; Wang et al. 2016), fish (Martins et al. 2006; Da Silva et al. 2012; Symonová et al. 2013, 2017), protists (Gong et al. 2013), insects (Cabral-de-Mello et al. 2010, 2011; Nguyen et al. 2010; Panzera et al. 2012; Palacios-Gimenez and Cabral-de-Mello 2015), bivalves (Pérez-García et al. 2014), and mammals (Sotero-Caio et al. 2015), and is regarded as strong evidence of rapid rearrangements over short time scales (Jiang and Gill 1994; Raskina et al. 2004, 2008). Mobilization of such multicopy gene families into heterochromatic regions is thought to be mediated through processes such as retrotransposon activity (Dimitri et al. 1997; Dimitri and Junakovic 1999; Symonová et al. 2013; de Bello Cioffi et al. 2015) and ectopic recombination (Nguyen et al. 2010).

Here we investigate patterns of rDNA CNV profiles in the *breve* group at two levels: the variation across specimens within species, and the variation among species. We focus our efforts on sequence-based evidence derived from low-coverage whole-genome sequencing data, but also validate sequence-based patterns using cytogenetic approaches. We survey the distribution of rDNA profile variation across the broader taxonomic group that contains the *breve* group (subgenus *Plataphus* of *Bembidion*). As part of our investigation in the *breve* group, we outline a simple approach to visualizing differences in the distribution of rDNA using copy number profiles generated by mapping reads to a reference and comparing the signatures resulting from copy number variation across specimens. Development of additional sequence-based approaches to detect variation in components of genomic architecture that can be easily and inexpensively measured from any specimen has potential to add clarifying signal to studies in species delimitation and genome evolution.

## Methods

### Overview

We investigated patterns of CNV within the ribosomal cistron across a framework of species recently delimited using evidence from molecular, morphological, and geographic data in Sproul and Maddison (2017). For each of the nine recognized *breve* group species (Fig. S1), we selected 3–8 specimens from across the species’s geographic range to test whether signatures observed in rDNA profiles were variable among, and stable within, putative species boundaries. We generated rDNA profiles by obtaining low-coverage whole-genome sequencing data and mapped reads for each specimen to a 14K-base outgroup reference sequence of the rDNA cistron of *Bembidion aeruginosum*. We chose *B. aeruginosum* as our phylogenetic studies (see below) indicate it is the sister group of the remaining *breve* group species. We conducted parameter sensitivity analysis for generating profiles, studied the effect of reference bias, compared profiles obtained from males and females, explored stability of profiles across varying read depth, tested whether profiles could be obtained from targeted sequencing workflows (i.e., hybrid capture), and searched patterns correlated with geography or phylogenetic patterns within species.

We used fluorescence *in situ* hybridization (FISH) to test the assumption that regions showing inflated copy number (CN) in rDNA profiles represent mobilization events in which fragments of rDNA have spread to new loci throughout the genome. We further validated patterns observed in rDNA profiles, and explored variation in repeats outside of rDNA, by conducting analysis of repetitive genomic elements using RepeatExplorer (Novák et al. 2010, 2013). We tested for broader taxonomic variation in rDNA profiles by generating profiles for 41 species of the subgenus *Plataphus*, the clade that contains the *breve* group. Our methods are explained in more detail below, and in Supplementary Materials.

### rDNAProfile Variation in the *breve* Species Group

We imported paired-end reads into CLC Genomic Workbench v9.5.3 (CLC Bio, referred to below as CLC GW), with failed reads removed during import. We trimmed and excluded adapter sequences from reads in CLC GW. We randomly down-sampled trimmed reads to 10 million per specimen, so that downstream analyses for all samples had a standardized number of input reads. We mapped trimmed reads to a ∼14K-base reference sequence of the rDNA cistron obtained from a *de novo* assembly of reads from *Bembidion aeruginosum* using the ‘Map Reads to Reference’ tool in CLC GW (match score=3, mismatch=4, insertion cost=3, deletion cost=3, length fraction=0.85, similarity fraction=0.85). We chose read mapping parameters following a sensitivity analysis in which we repeated read mapping across a range of parameter settings using four representative samples. Additional methods used for the parameter sensitivity analysis, for screening mapped reads for contaminants and assembly artifacts, and for obtaining the rDNA reference sequence are provided in Supplementary Materials, Figures S2–3, and Table S3. Following read mapping, we removed duplicate mapped reads in CLC GW.

We visualized the pattern of coverage depth resulting from read mapping by generating graphs of read pileups in CLC GW. These graphs of coverage depth form the initial visual component of rDNA profiles presented here (Fig. 1). We enhanced these graphs by converting read depth to copy number (see Supplementary Materials), and applying a color ramp in Illustrator to indicate the magnitude of copy number differences within the profile. We applied the color ramp such that the color of all rDNA profiles shown here indicates copy number relative to the maximum value observed (206,239 copies, *B. lividulum* 5013) throughout all of the profiles (Figs. 1–2).

**Figure 1.**
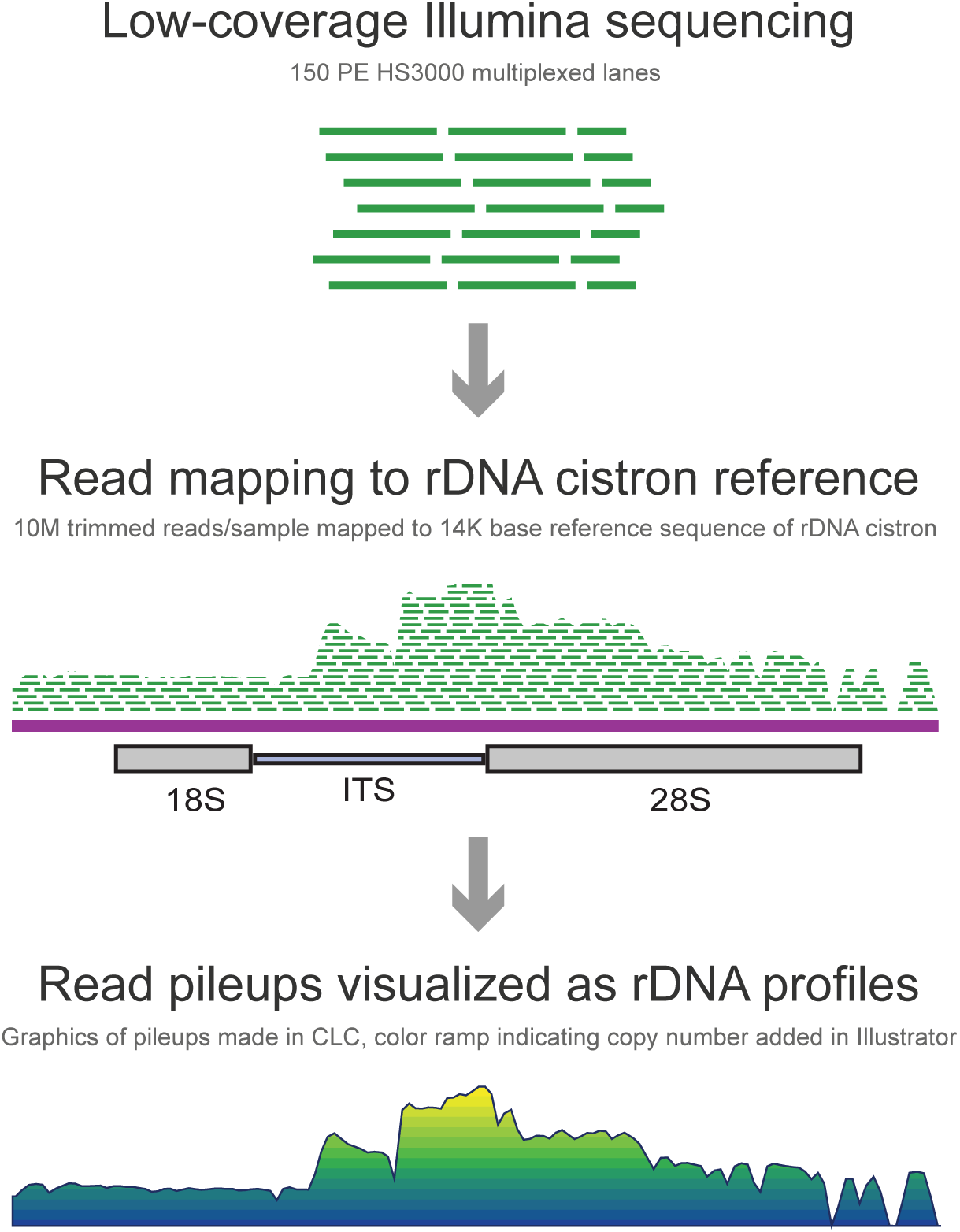
Flowchart illustrating the steps used to generate rDNA profiles from short-read sequencing data.

**Figure 2.**
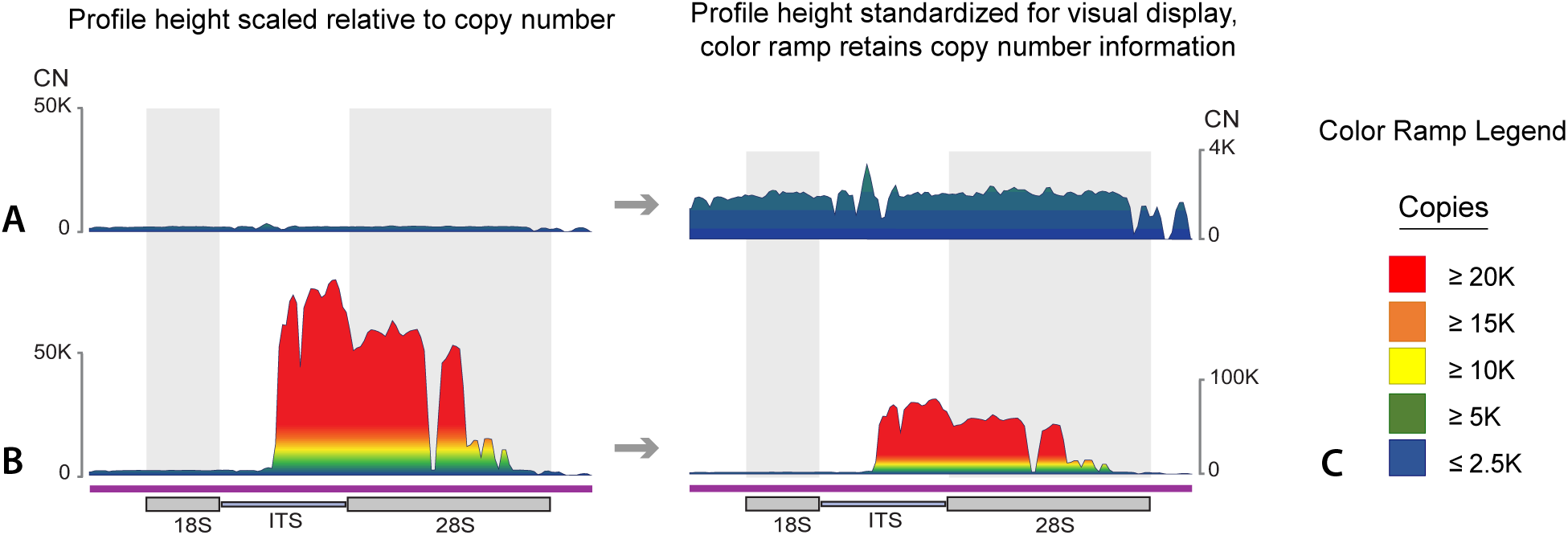
A comparison of rDNA profiles shown with relative vs. fixed scales on the y-axis for two specimens: (A) *Bembidion laxatum* (5086), and (B) *B. lividulum* (3486). Profiles on the left are scaled relative to 50,000 copies, and the same profiles on the right constrained to the same maximum height. We use the latter scaling strategy throughout the paper to simplify visual display of rDNA profiles. (C) To emphasize differences in CN that are less apparent in profiles that are scaled to a uniform height, we applied a standardized color ramp to all rDNA profiles included in this study such that any region with >20K copies = red, >15K copies = orange, >10K copies = yellow, >5K copies = green, and <2.5K copies = blue.

We note that the approach used to generate rDNA CNV profiles described above relies on CLC GW, which is commercial software with proprietary algorithms. In Supplementary Materials we outline an alternative workflow using freely available, open algorithms for generating CNV profiles using freely available software tools Bowtie2 v2.3.4.2 (Langmead and Salzberg 2012), SAMtools v1.9 (Li et al. 2009) and R v3.5.1 (R Core Team 2013).

### Evaluating rDNA profile variation within and among species

We mapped rDNA profiles obtained for all *breve* group specimens onto the tree used to infer species boundaries by Sproul and Maddison (2017) in order to determine the extent of rDNA profile variation among species, and whether distinctive features in rDNA profiles (e.g., position of regions showing CN inflation, and the magnitude of inflation in those regions) within a species showed stable signatures across individuals sampled from diverse geographic localities.

We conducted within- and between-species analysis of rDNA profile shape by testing for correlation in coverage depth patterns across the rDNA cistron for all *breve* group specimens. Using the BAM files from which we generated rDNA profiles, we calculated coverage depth at each position across the rDNA cistron for each sample using the “depth” command in SAMtools v1.9 (59). In this way, we converted each profile into ∼14K point depth estimates, one at each position along the reference sequence to which reads were mapped. We then calculated Spearman’s rank correlation coefficient (or Spearman’s rho, denoted ‘*ρ*’) for pairwise comparisons of all *breve* group specimens. Spearman’s rho is a nonparametric measure of rank correlation, which in this case is measuring the degree of similarity in the variable of coverage depth for each rank (or position) across the rDNA cistron between two samples. We calculated Spearman’s rho and generated a histogram of rho values for all pairwise comparisons in R v3.5.1 (60).

We classified rDNA profiles based on the presence of CN inflation within the rDNA cistron as follows: “high” CN inflation (profiles in which maximum CN ≥ 20-fold higher than baseline CN); “moderate” CN inflation (maximum CN ≥ 10–19.99-fold higher than baseline CN); “low” CN inflation (maximum CN ≥ 3–9.99-fold higher than baseline CN); and “lacks” CN inflation (maximum CN < 3-fold higher than baseline CN).

### Cytogenetic Mapping of Ribosomal DNA

We performed FISH experiments with three *breve* group species, using FISH probes to target regions of 18S and 28S rDNA that vary in copy number within and among species. We performed tissue dissection and fixation following Larracuente and Ferree (Larracuente and Ferree 2015), and conducted FISH using protocols that combined steps from Larracuente (Larracuente 2017) and Symonová *et al*. (Symonová et al. 2015). We confirmed results using multiple probe synthesis and post-hybridization wash strategies, and with multiple fluorophores. Additional details of FISH methods are provided in Supplementary Materials and Table S4.

## Results

### rDNA Profile Variation in the *breve* Species Group

An overview of methods used to generate and display rDNA profiles shown here is provided in Figures 1 and 2.

Ribosomal DNA CNV profiles generated from the *breve* species group showed species-specific signatures of variation across the group (Fig. S3). Five of nine species showed unique regions with inflated CN (i.e., 3–100+ fold CN increase) relative to the rest of the rDNA cistron (Fig. 3, Table S2). Two species (*Bembidion lividulum* and *B. breve*) showed high CN inflation, two species (*B. geopearlis*, *and B. testatum*) showed moderate CN inflation, and one species (*B. saturatum*) showed low CN inflation (Fig. S3, Table S2). Although profiles for the remaining four species (*B. ampliatum*, *B*. *laxatum*, *B. oromaia* and, *B. vulcanix*) lacked CN inflation, species-specific signatures were still evident in the pattern of regions with reduced read mapping coverage (e.g., the position of valleys in the rDNA profiles), as well as minor peaks (e.g., peaks less than 3-fold higher than the baseline CN). This variation is primarily due to species-specific patterns of sequence divergence and indel location relative to the reference sequence (Fig. S3).

**Figure 3.**
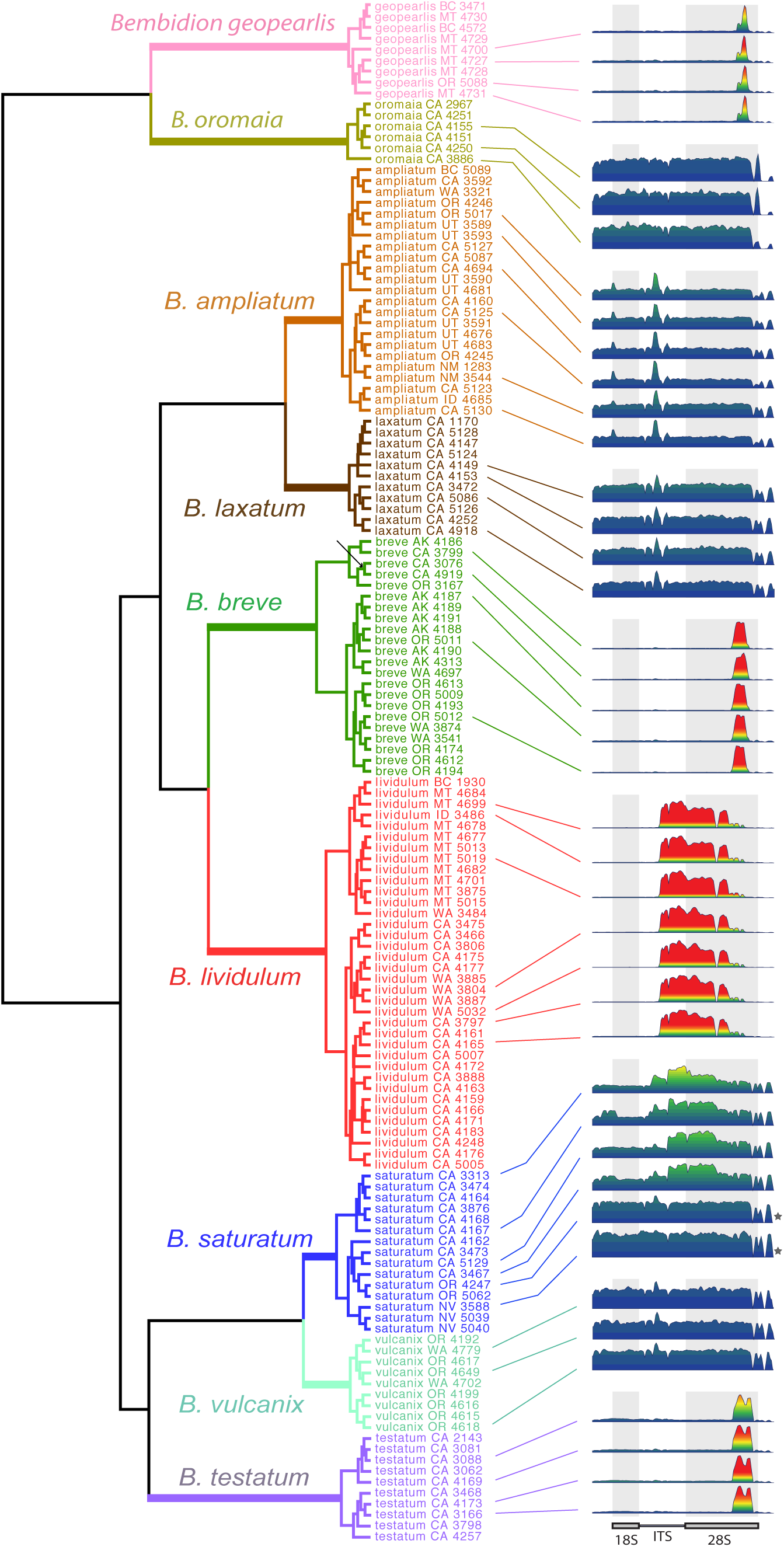
The tree used to infer species boundaries of the *breve* species group (adapted from Sproul and Maddison (2017), Fig. 7) with rDNA profiles. Terminal taxa are colored by inferred species. rDNA profiles for several specimens of each species are shown to the right of the terminals. One to two profiles for some species (e.g., *Bembidion lividulum* and *B. ampliatum*) were excluded to facilitate visual display; however, all profiles not shown corroborate patterns evident in the figure. Profiles for two morphologically distinct specimens suspected of belonging to cryptic lineages in Sproul and Maddison (2017) are indicated by gray stars. All profiles generated are shown in Figs. S5–S13. Branch length is proportional to relative divergence with scale bars indicating 0.01 units.

Species-specific signatures observed in rDNA profiles were highly stable across multiple individuals of each species sampled from various geographic localities (Fig. 3 and S5–S22). In particular, the boundaries (i.e., the exact position in the rDNA cistron) of regions showing inflated CN were stable within species, and across varying read mapping parameters (Figs. 3 and S2). Within regions showing CN inflation, maximum CN was somewhat variable within species, most notably in *B. lividulum* and *B. testatum,* which both showed greater than 3.5-fold variation in maximum CN across specimens (Tables S2 and S9; Figs. S5 and S12).

Spearman’s rho showed very strong correlation of CNV profile shape, as measured by read depth patterns across the rDNA cistron, for within-species comparisons (average *ρ* of all within-species comparisons=0.966, SD=0.032; range of average *ρ*=0.933–0.998) (Figs. 4 and S23, Table S9). The strength of correlation for between-species comparisons varied widely (average *ρ* of all between-species comparison =0.347, SD=0.306). Correlation in comparisons for which one or both species showed CN inflation was generally low (average *ρ* =0.263, SD=0.251), and moderate to strong in specimens of species which both lacked CN inflation (average *ρ* =0.7883, SD=0.035, range *ρ* =0.674–0.873), but not so strong as between-species correlation (Figs. 4 and S23, Table S9).

Pairwise comparisons that included a rDNA profile obtained from a female specimen consistently showed rho values that were as high or higher than male-male pairwise comparisons (Figs. S5–S13, Table S10). Similarly, comparisons in which one profile was obtained from our hybrid capture workflow had rho values within the range of variation seen in profiles generated using our standard approach (Figs. S5–S13, Table S10). Profiles generated from the same specimen using 10 million, 5 million, and 1 million reads were all nearly identical (Fig. S23).

The average total fraction of reads mapping to the 14K-base outgroup reference sequence ranged from an average of 1.07% (SD=0.2%, n=7) in *B. ampliatum* to 14.7% (SD=2.8%, n=8) in *B. lividulum* (Table S2).

Ribosomal DNA profiles for the two specimens of unknown taxonomic status that are currently classified under *B. saturatum* lacked a broad region of CN inflation present in all *B. saturatum* specimens, and accordingly showed only moderate correlation in profile shape with *B*. *saturatum* specimens (average *ρ* = 0.580, SD=0.043) (Fig. 4).

**Figure 4.**
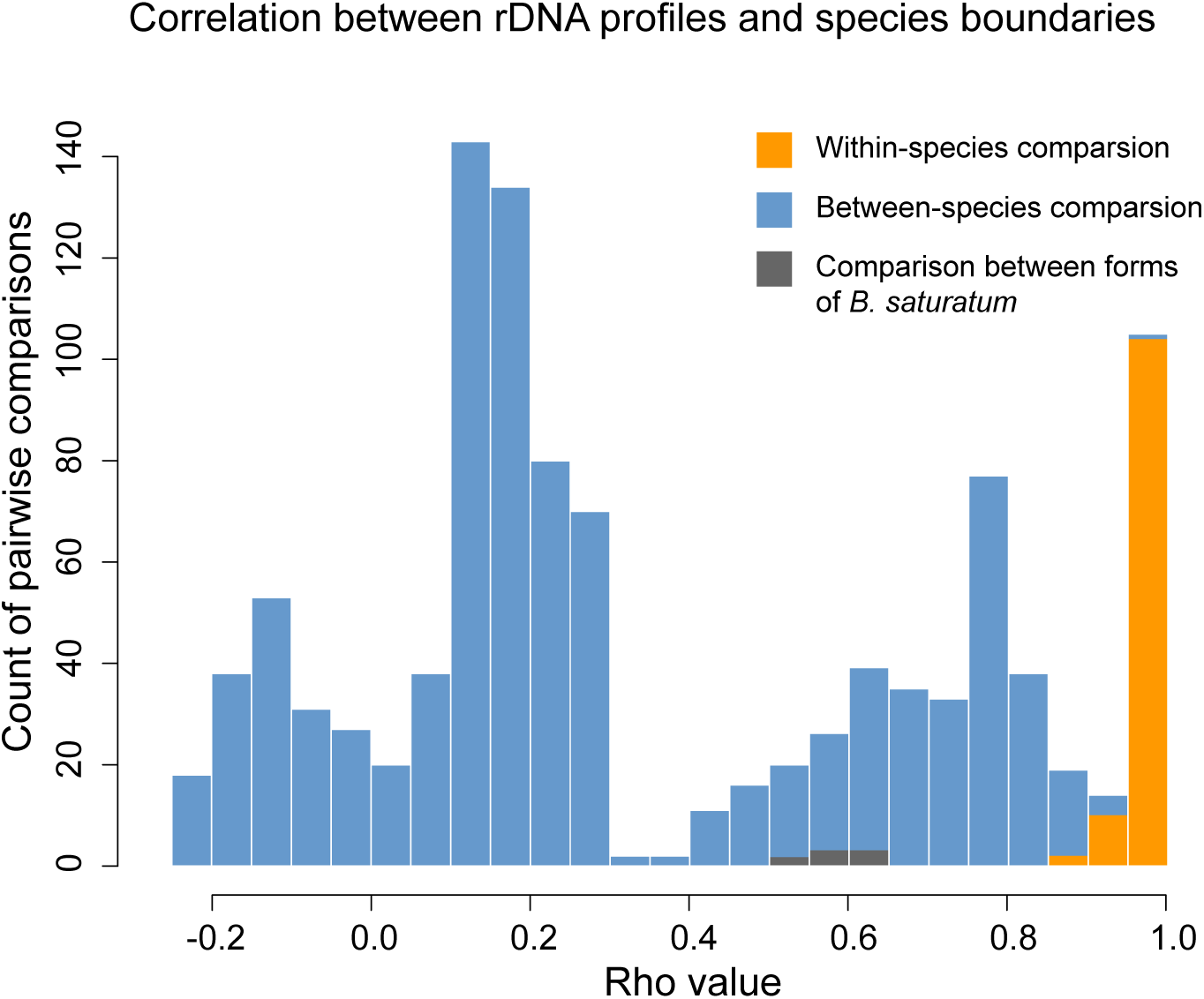
A histogram of rho values summarizing the results of the correlation analysis between rDNA profiles of specimens, with comparisons within and between species indicated.

Among specimens of *B. lividulum* we noted minor variation in profile shape (i.e., copy number) within the inflated region; this variation was consistent with phylogenetic position and geographic locality of the specimens sampled (Figs. 5, S5, and S14). We noted minor variation in other species (e.g., *B. saturatum*), but this did not show obvious correlation with phylogenetic or geographic patterns (Figs. S6 and S15).

### Cytogenetic Mapping of Ribosomal DNA

Patterns observed in fluorescence *in-situ* hybridization (FISH) experiments corroborated our hypothesis that copy number inflation of specific regions within the rDNA cistron is due to mobilization of rDNA, and can represent substantial variation in location and abundance of rDNA-like repeats across the genome. In all species, hybridization with probes targeting rDNA regions lacking CN inflation (18S in *B. lividulum*, 18S and 28S in *B. vulcanix*, and 28S in *B. testatum*) produced two strong FISH signals, whereas hybridization with probes designed in CN inflated regions (marked 28S inflation in *B. lividulum*, and slight 18S inflation in *B. testatum*) showed more than two FISH signals (4–5 loci in *B. testatum*, and many loci in *B. lividulum*) (Figs. 6, 7 and S25). In *B. lividulum*, sufficient tissue and replicate squashes were available to confirm the distribution of FISH signals on condensed, well-spread chromosomes. Uninflated 18S rDNA mapped to two chromosomes, whereas markedly inflated 28S rDNA showed FISH signals on portions of all 24 chromosomes (Fig. 7). Based on known position of euchromatin/heterochromatin boundaries in *Bembidion* chromosomes in meiotic chromosomes (Maddison 1986), the pattern of FISH signals we observe suggests that much of the mobilized 28S rDNA-like sequences are concentrated in heterochromatic regions of chromosomes and frequently absent on euchromatic tails (Fig. S26).

The fluorescence patterns seen in *B. testatum* chromosomes were the same whether the sequences of the 28S probes matched those of *B. lividulum* or *B. testatum*. Similarly, the fluorescence patterns seen in *B. lividulum* chromosomes were the same whether the sequences of the 28S probes matched those of *B*. *lividulum* or *B*. *testatum*.

### Cluster Analysis of Repetitive DNA

Cluster analysis of genomic repeats corroborated general patterns observed in rDNA profiles within the *breve* group. *Bembidion lividulum* had an average of 21 clusters containing rDNA hits whereas none of the species that lacked CN inflation had more than four clusters with rDNA hits. Beyond rDNA, the composition of major repeat categories was somewhat variable below the species level, though some species-specific trends were evident (Figs. S5–S13). For example, clusters of simple repeats (e.g., satellite DNA) were consistently more abundant in *B. ampliatum* specimens than in other species (Fig. S7), whereas clusters of Class I transposable elements (TEs) were notably abundant in *B. breve* (Fig. S8). Female specimens of all species lacked (or had reduced) Class II TEs compared to male specimens of the same species (Figs. S5– S13). *B. lividulum* showed variation in major repeat categories and superfamilies of Class I and Class II TEs that followed geographic and phylogenetic patterns similar to rDNA profiles of the same specimens (Fig 5).

**Figure 5.**
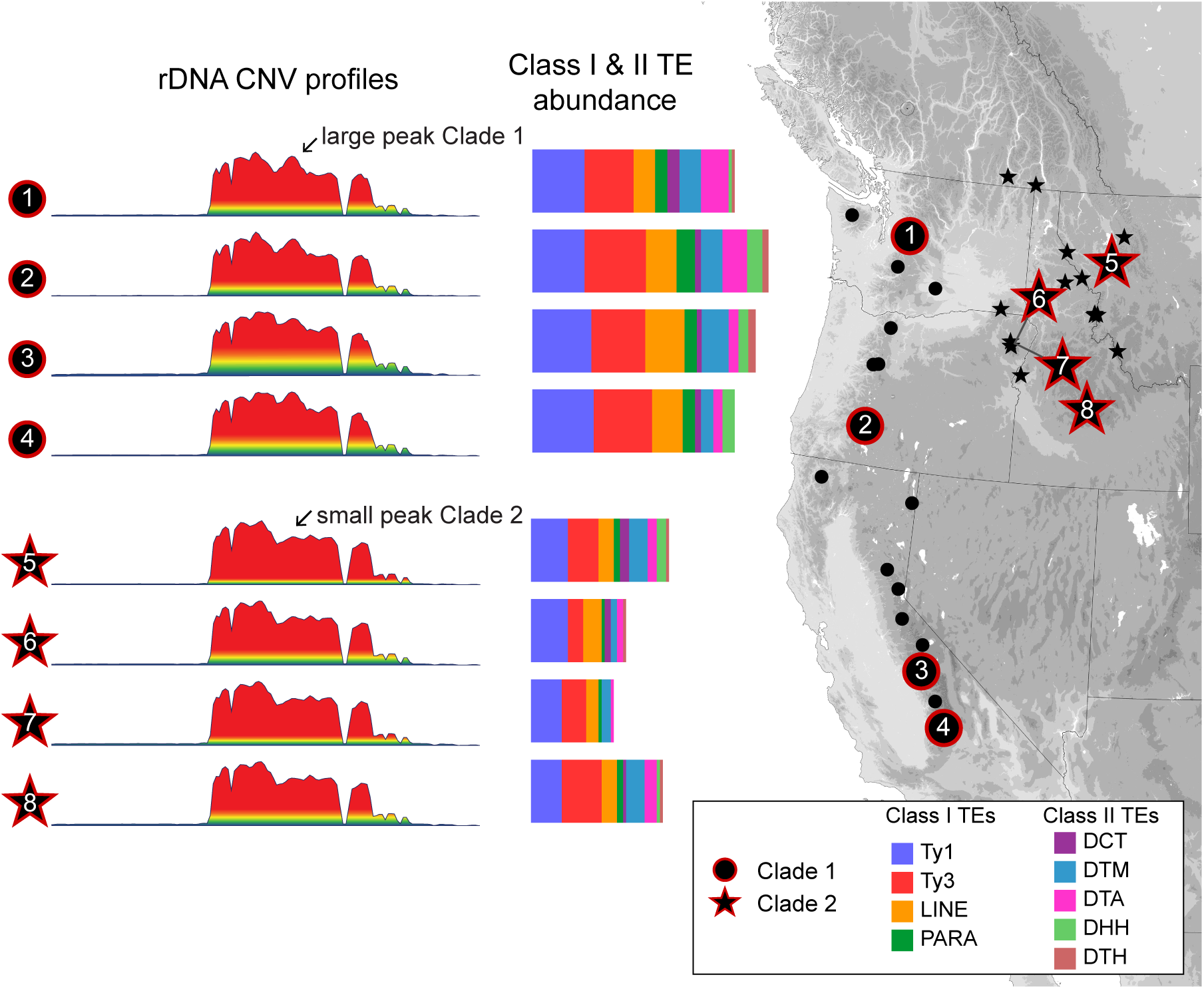
Summary of data obtained from *Bembidion lividulum* specimens including rDNA profiles, repeat content of Class I & II TE superfamilies, and a map of western North America showing sampling localities of the sampled *breve* group specimens. Localities with specimens belonging to Clade 1 are shown by circles, while localities with specimens belonging to Clade 2 are shown by stars. Circles and stars outlined in red indicate localities from which we obtained rDNA profiles.

### *rDNA Profile Variation across* Bembidion (Plataphus)

We found evidence in rDNA profiles that rDNA mobilization is widely distributed across the subgenus *Plataphus* (Fig. 8). Seventeen of 41 non-*breve* group *Plataphus* specimens showed CN inflation, with four species showing high inflation, seven with moderate inflation, and six with low inflation (Fig. 8, Table S8). Five of the ten major clades in the subgenus had one or more species with CN inflation in rDNA profiles (Fig. 8).

The distribution of rDNA profile variation across the subgenus *Plataphus* showed two general phylogenetic patterns. We observed strongly discordant profile signatures between sister groups, suggesting rapid divergence of CNV profiles among species that show little divergence in other genomic regions (e.g., sequence divergence of single-copy genes). There were eight instances in which rDNA profiles from a given species showed rDNA regions with greater than 10-fold increase in maximum copy number relative to the same region in their sister taxon (or one or more species in their sister group) (Fig. 8, Table S2).

In contrast, in some regions of the phylogeny, small clades shared similar patterns of inflation. For example, all species sampled from the *planiusculum* group showed inflation at the same ITS region, though the degree of inflation varied across species in the group, and two species showed additional regions of CN inflation in 18S (Fig. 8). In addition, three species within the *curtulatum* group showed inflation within a conserved region of 28S, though the magnitude and pattern of inflation across that region varied among species (Fig. 8). We did not observe cases in which signatures of inflation persisted across larger, older clades.

## Discussion

Comparative study of genome architecture across individuals and species has potential to illuminate new mechanisms underlying the complexity and diversity of life (Lynch and Walsh 2007), yet mapping of whole genome architecture in most groups remains a technical and financial challenge. In this study, rDNA CNV profiles provide an example of how a simple measure of one component of genome architecture can offer clarifying signal to evolutionary studies. In the *breve* group, rDNA profiles provide clean signal that is highly correlated with species boundaries (Figs. 3-4) in a complex group of very similar species, for which a multi-year study of individual gene trees, multi-gene analyses, and morphological characters was previously required for delimiting species (Sproul and Maddison 2017).

Initially, we found the consistency of rDNA profile signatures within species to be surprising given that repetitive DNA is known to be dynamic even at the sub-population level (West et al. 2014). In fact, aspects of our analyses supported the dynamic nature of repeats below the species level. For example, our cluster analysis revealed substantial within-species variation of repetitive DNA categories and abundance of Class I & II TEs, such that only general trends showed species-specific signatures (Figs. 5 and S5– S13). In addition, the maximum copy number of inflated rDNA regions showed greater than 3-fold variation within species (Table S2), likely due to expansion and contraction of mobilized arrays as a result of unequal exchange (Szostak and Wu 1980; Charlesworth et al. 1994; Eickbush and Eickbush 2007). Despite this within-species variation in absolute copy number, our results demonstrate that by using a common reference sequence to generate a copy number profile of a specific repeat, it is possible to cut through the noise caused by the dynamic nature of repeats and view a condensed summary of evolutionary events (e.g., sequence divergence, indel accumulation, and repeat mobilization) that provides a stable signal informative to studies at the species level. In some cases, these events have minimal impact on the structure of the genome as a whole (e.g., sequence divergence, indel accumulation, or minor mobilization events as in Fig. S3). But in other cases, they represent the re-patterning of a major repetitive component of the genome (e.g., the degree of rDNA mobilization in *B. lividulum*, *B. breve, B. haruspex*, and *B.* sp.nr. *curtulatum* “Idaho”) (Figs. 6–8, and S26), and a simple measure of that component can add strong evidence regarding species boundaries that is not evident in more commonly considered data sources such as gene-tree analysis of individual genes. Our findings add to the growing body of literature that uses low-coverage sequence data and novel approaches to extract signal of genome-scale variation (West et al. 2014; Dodsworth et al. 2015; Denver et al. 2016; Lower et al. 2017), and highlights repeats as an underdeveloped source of signal for evolutionary studies (Dodsworth et al. 2015).

**Figure 6.**
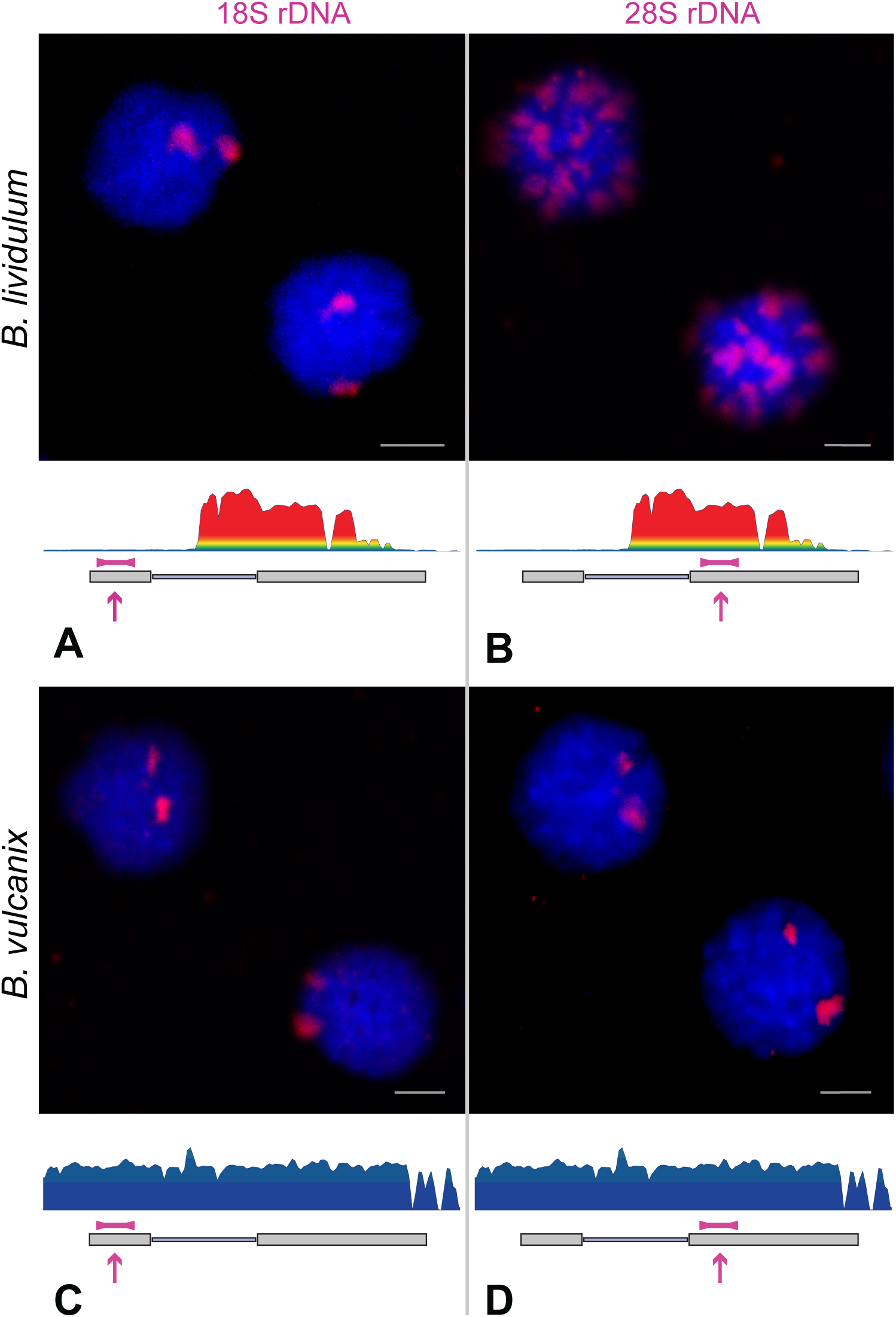
FISH signals obtained by cytogenetic mapping of rDNA in *Bembidion lividulum* and *B. vulcanix*. (A) FISH signals resulting from hybridization of 18S probes to *B. lividulum* nuclei; (B) FISH signals resulting from hybridization of 28S probes to *B. lividulum* nuclei; (C) FISH signals resulting from hybridization of 18S probes to *B. vulcanix* nuclei; (D) FISH signals resulting from hybridization of 28S probes to *B. vulcanix* nuclei. Ribosomal DNA profiles for *B. lividulum* and *B. vulcanix* are shown below their respective FISH images with the position of either 18S or 28S FISH probes indicated by pink boxes and arrows.

**Figure 7.**
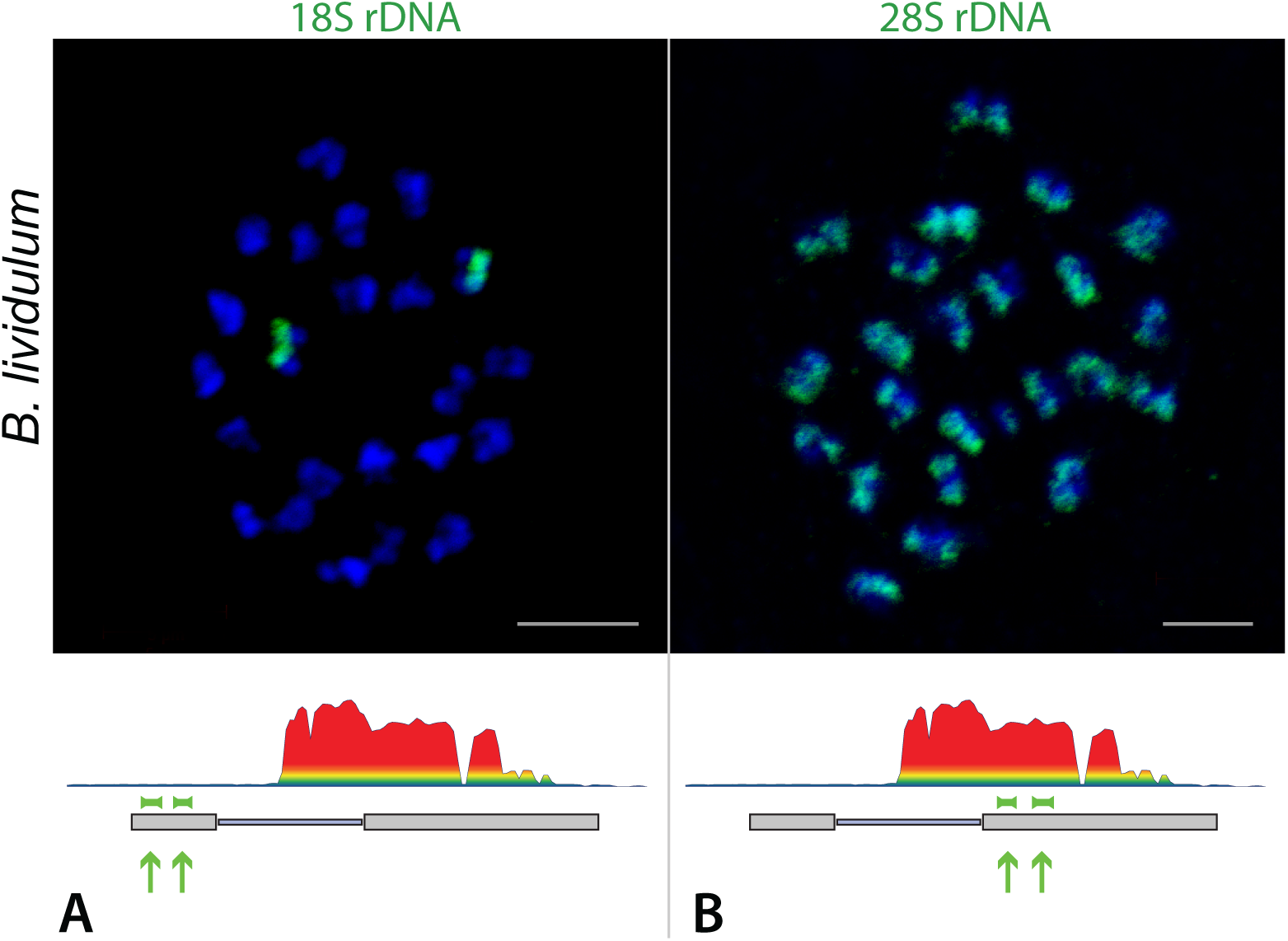
FISH signals obtained by cytogenetic mapping of rDNA in *Bembidion lividulum*. (A) FISH signals resulting from hybridization of 18S probes to condensed chromosomes (B) FISH signals resulting from hybridization of 28S probes to condensed chromosomes. Ribosomal DNA profiles for *B. lividulum* are shown below FISH images with the position of either 18S or 28S probes indicated by green boxes and arrows.

**Figure 8.**
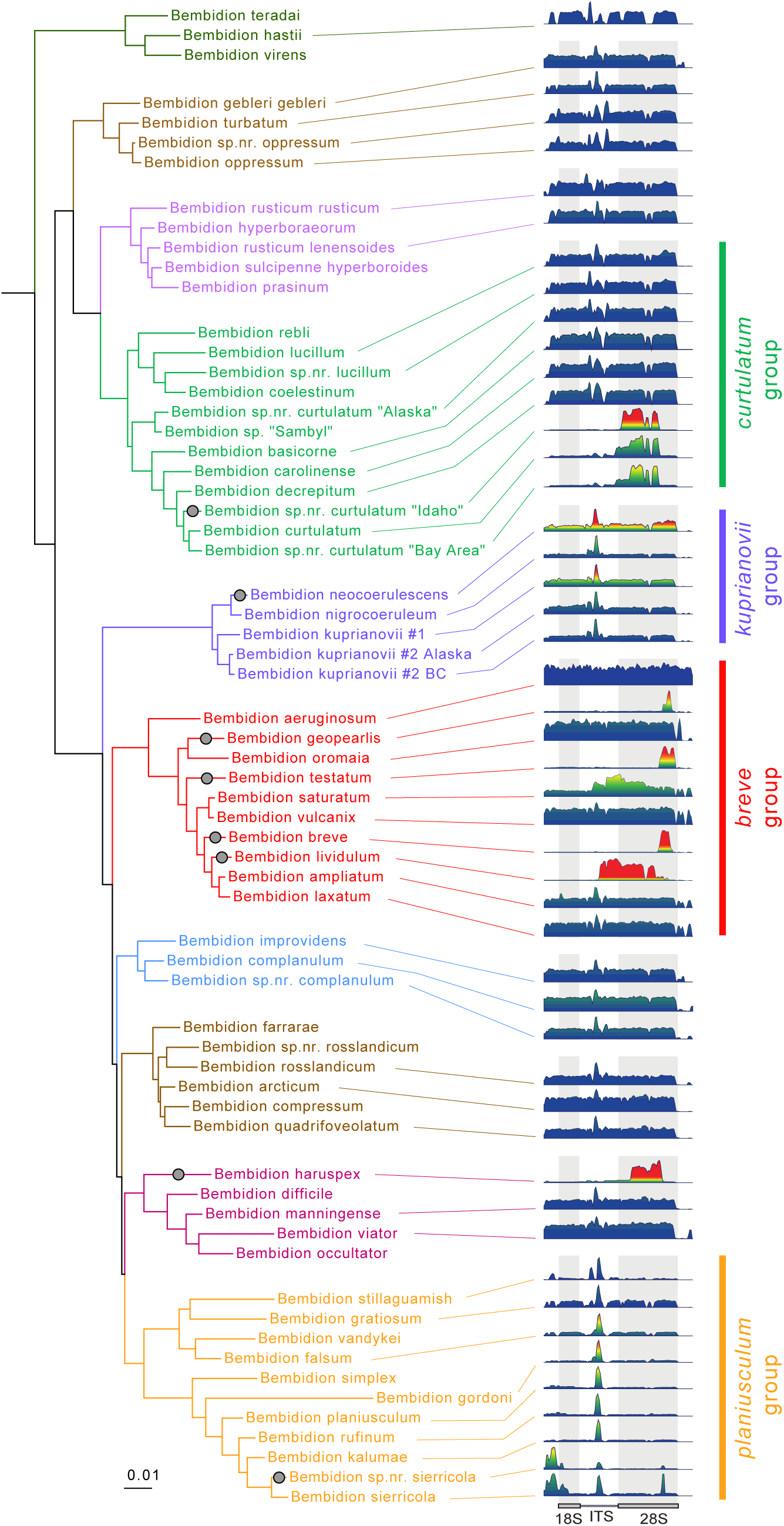
Maximum likelihood tree of *Bembidion* subgenus *Plataphus*, the subgenus containing the *breve* species group, with rDNA profiles for many species. The species groups discussed in the text are indicated with colored bars and text to the right of rDNA profiles. Species with regions in rDNA profiles that differ >10-fold in copy number relative to the same region in a sister species, or one or more species in their sister group, are indicated with a gray circle.

Cytogenetic mapping of rDNA demonstrated that patterns of CN inflation observed in rDNA profiles correspond to re-patterning of the abundance and relative position of rDNA-like sequences throughout the genome (Figs. 6, 7 S25–26). This finding validates our hypothesis that rDNA CNV profiles summarize one aspect of genomic architecture in that they can identify repeat regions that contribute to variation in the relative linear position of repeat arrays across chromosomes. Linear patterning of genomic components is central to determining DNA-DNA and DNA-protein interactions in the nucleus, and empirical studies have demonstrated that shifts in the abundance and position of blocks of repeats alter patterns of chromatin formation (e.g., heterochromatin/euchromatin boundaries), gene expression, and phenotypes (Wallrath and Elgin 1995; Lemos et al. 2010; Elgin and Reuter 2013), and are hypothesized to be a mechanism that underlies reproductive isolation of some recently diverged lineages (Ferree and Barbash 2009; Feliciello et al. 2014; Hall et al. 2016). The relative importance that shifts in one repeat class might have on entire genomic architecture is expected to vary widely among repeats and organisms, and cannot be inferred from a CNV profile. However, surveys of such variation can serve to identify new model systems for study, and efficiently direct efforts of more costly approaches to investigate the role of repeat architecture on genome evolution and speciation.

Our FISH analysis also establishes a link between rDNA profiles and the results of many cytogenetic studies across the tree of life that document rDNA movement as a driver of genome evolution over short time scales (Raskina et al. 2008; Panzera et al. 2012; Gong et al. 2013; Symonová et al. 2013; Sember et al. 2015), and demonstrates a low-cost, sequence-based measure to visualize this long-studied source of variation. Although our sequence-based approach lacks fine-scale details (such as locations within chromosomes) provided by cytogenetic mapping techniques, it has the advantage that it can be applied to any specimen for which DNA sequences can be obtained, including specimens with old and ancient DNA (Fig. S27). Because DNA sequencing projects can be designed for increasingly high throughput, CNV profiles of rDNA, or profiles of other repeats, have excellent potential as a tool for identifying genomic components that contribute to genome-scale variation across groups, including in groups that lack pre-existing genomic resources (e.g., annotated reference genomes).

Phylogenetic sampling of rDNA profiles across 50 species in the subgenus *Plataphus* showed that rDNA mobilization events have been relatively common in the recent evolutionary history of the group (Figs. 3 and 8). We did not detect rDNA mobilization events deeper in the phylogeny of *Plataphus*, but we expect older events would be undetectable by our methods. If earlier mobilizations did occur, they would now be invisible to our read-mapping approach if mobilized rDNA escapes concerted evolution with functional rDNA clusters, and diverges sufficiently. This observation is consistent with cytogenetic studies on the distribution of rDNA that include broad taxonomic sampling (Nguyen et al. 2010; Cabral-de-Mello et al. 2011; Sember et al. 2015; Wang et al. 2016). Thus, our inferred number of mobilization events is a lower bound, and the true number of mobilization events in the history of *Plataphus* could well be higher. However, the fact that regions showing CN inflation are highly stable across individuals within species (Fig. 3) suggests that rDNA mobilization events are sufficiently rare as to allow for fixation of the signature across individuals within species (likely facilitated through concerted evolution among mobilized clusters), but sufficiently common as to frequently show different patterns between closely related species. Our finding eight instances of sister-group pairs that differ strongly in rDNA profile features indicates that genomic differentiation through re-patterning of rDNA-like sequences has occurred frequently and rapidly within *Plataphus*. Such genomic restructuring of rDNA-like sequences has been hypothesized in other taxa to drive speciation (Raskina et al. 2004; Symonová et al. 2013), and this could have played a role in the diversification of *Bembidion* subgenus *Plataphus*. The highly stable nature of rDNA profile shape within species, together with the pattern of dramatic variation in rDNA profiles between species, provides a strong visual illustration of the paradox that although rDNA is one of the most highly conserved fractions of the eukaryotic genome, it can be simultaneously a hypervariable driver of genome evolution (Raskina et al. 2008; Gibbons et al. 2014; Malone 2015).

## Acknowledgements

We thank Barbara Taylor, Amanda Larracuente, Radka Symonová, and James Strother for their advice in optimizing FISH protocols. We thank Anne-Marie Girard-Pohjanpelto for help with fluorescence imaging. We are grateful to Danielle Mendez for her help with library quantitation. We thank Elizabeth Sproul, Steven Wainwright, Aaron Liston, and Amanda Larracuente for editorial input on the writing. We thank Olivia Boyd, Ching-Ho Chang, William Cresko, Dee Denver, Michael Freitag, Tiffany Garcia, Antonio Gomez, Lucas Hemmer, Kojun Kanda, David Kavanaugh, Danielle Khost, Amanda Larracuente, Aaron Liston, David Lytle, Beatriz Navarro-Domínguez, Eli Meyer, Wendy Moore, James Pflug, José Serrano, Xiaolu Wei, and Kipling Will for discussions that improved the scope of the work presented here. We thank Elizabeth Sproul, George Sproul, Pearl Sproul, Janet Reed, and Greg Reed for their help collecting specimens used for FISH. Newly reported sequence read files are deposited in NCBI Sequence Read Archive (Accessions SAMN10860544– SAMN10860633). PCR-based sequence data are deposited in GenBank (Accessions MK461576 – MK461828). This work was supported by the National Science Foundation (grant numbers DEB-1702080 to JSS and DRM, DEB-1258220 to DRM, BIO-1812279 to JSS); and the Harold E. and Leona M. Rice Endowment Fund at Oregon State University.

