## Supplemental Materials for "Repetitive DNA profiles Reveal Evidence of Rapid Genome Evolution and Reflect Species Boundaries in Ground Beetles"

**This PDF file includes:**

Supplemental Methods
References for supplemental reference citations
Supplemental figure captions
Figs. S1 to S27
Tables S1 to S9

### Supplemental Methods

**Taxon sampling and DNA extraction for *breve* species group specimens**

We generated low-coverage whole-genome sequence data from 41 *breve* group specimens selected from the taxon sampling of Sproul and Maddison (2017a)(Sproul and Maddison 2017a), with 3–8 specimens chosen per species (Table S1). We obtained Illumina reads for six additional specimens (3593, 4149, 4165, 4245, 4918, and 5032) from Sequence Read Archive (SRA SRR5514451–SSR5514456). Reads obtained from SRA were generated previously by the authors using the same library preparation and sequencing protocols as newly sequenced specimens (Sproul and Maddison 2017a, 2017b). For one species, *B. saturatum*, we added to our sampling two individuals of unclear taxonomic status. Sproul and Maddison (2017a) noted that individuals from two disjunct populations of *B. saturatum* showed distinct morphological forms; however, because gene trees showed limited support for their distinctiveness, and because only a few specimens from disjunct geographic localities were available, the authors deferred taxonomic action until more data could be gathered. DNA extraction protocols are provided in Sproul and Maddison (2017a).

**Library preparation and sequencing**

We prepared DNA extractions for Illumina sequencing by sonicating genomic DNA using a Diagenode Bioruptor Pico Sonicator, and prepared dual-indexed libraries using NEBNext DNA Ultra II Library Prep Kits (New England Biolabs) following the manufacturer's recommended protocol. We attempted to maximize library evenness by standardizing input DNA quantities between 10–50 ng (except for five samples, three having less than 10 ng of available DNA, and two for which maximizing input was desirable due to their role in other projects), and by using a consistent number of library amplification cycles for a given amount of input (Table S2). The libraries were then pooled and sequenced on one of four 150 base paired-end lanes on an Illumina HiSeq 3000 maintained by the Oregon State University Center for Genome Research and Biocomputing. We allocated as much as 1/4 and as little as 1/52 of a lane per sample. Lane pairings are provided in Table S2. Following sequencing, demultiplexing of Illumina reads was performed using CASAVA v1.8 (Illumina). Reads are deposited in Sequence Read Archive with accession numbers SAMN10860544–SAMN10860633.

**Obtaining rDNA reference sequence**

We obtained the rDNA reference sequence through *de novo* assembly of *Bembidion aeruginosum* reads in CLC GW with default settings. We created a BLAST database of the resulting contigs and used both 18S rRNA and 28S rRNA gene sequences obtained from *B. aeruginosum* (via PCR and Sanger sequencing) as query sequences in a BLAST search against the database of contigs. The best scoring hit for both queries was a single contig ~14k bases in length. We annotated boundaries of rRNA genes on the contig using RNAmmer 1.2 Server (Lagesen et al. 2007). We did not find 5.8S in our *B. aeruginosum* assemblies even after using multiple assembly strategies; the current reference sequence represents 45S rDNA without 5.8S. The lack of 5.8S may contribute to read mapping artifacts observed in ITS regions (Fig. S3).

Using a single reference sequence for read mapping of all study specimens was desirable as it allowed for comparing variation in rDNA profiles from a fixed point of reference. An alternative approach would be to generate rDNA profiles by obtaining a reference sequence of the rDNA cistron for each species individually, and mapping reads to that reference. This approach would have the advantage of bypassing artifacts associated with using a non-conspecific reference (e.g., underestimates of coverage in regions with high sequence divergence) (Fig. S3); however, it has the disadvantage of preventing the direct visual and statistical comparison of variable regions across specimens due to indels (that are common in rRNA genes and spacer regions), which can potentially result in variable reference sequence length across specimens. In addition, we found this approach to be time costly given that *de novo* assembly of the whole rDNA cistron was extremely difficult for any specimens that showed CN inflation within the rDNA cistron, in which inflated regions show very poor contiguity. In *Bembidion lividulum*, for example, we repeated *de novo* assembly after systematically down sampling reads from 10 million to 100,000 reads, and using multiple assembly algorithms; however, the assembly of the multi-thousand base inflated region of ITS and 28S remained broken into dozens of very short (e.g., 150-400 base) contigs. Thus, for both efficiency and comparative value, we found the use of a single reference to be the best approach to comparing rDNA profile patterns. Had our sampling extended to further taxonomic breadth, use of additional references would have likely been necessary due to increasing sequence divergence among taxa.

**Estimating copy number from coverage depth**

We converted coverage depth to copy number by dividing coverage depth values at a region of interest (e.g., maximum coverage depth) by the average coverage depth of 67 putatively single-copy nuclear protein-coding genes (Regier et al. 2008), which we mapped from the same set of reads (Sproul and Maddison 2017a). We estimated baseline copy number by calculating average coverage depth across a region of 18S (positions 5000–5500 in the reference sequence) which lacked copy number inflation for all species measured in this study (with one exception addressed below), and divided the average baseline coverage depth by the average coverage depth of the single-copy genes mentioned previously. The single specimen in the study that showed copy number inflation in the chosen 18S region was *Bembidion testatum*. For this specimen, we used a 500-base region of 28S lacking copy number inflation to calculate the baseline copy number.

**Effect of methodological approaches on rDNA profiles**

We studied the effect of reference bias on rDNA profiles by mapping reads to the standard reference of *B. aeruginosum* and comparing that profile to one produced using an alternative approach, in which the reads were mapped to a reference sequence obtained from *de novo* assembly of reads from the same species being mapped (Fig. S3).

We conducted a read mapping parameter sensitivity analysis by selecting four specimens, two from the *breve* group (*B. lividulum* DNA3486 and *B. breve* DNA4187) and two other *Plataphus* (*B. gordoni* DNA2358 and *B. sp.nr.curtulatum "*Idaho" DNA3613), which were chosen to represent a diversity of rDNA profile shapes. We repeated read mapping nine times for each specimen across varying stringency of parameter values (i.e., match score, mismatch score, insertion cost, deletion cost, length fraction, and similarity fraction) as listed in Table S3. We visualized rDNA profiles for all read mapping trials. Read mapping parameters were judged to be ideal if they produced a profile that had a stable shape compared to profiles produced using parameter settings of one level higher stringency, and one level lesser stringency for at least three of four taxa studied. An example of rDNA profiles resulting from the parameter sensitivity analysis is provided in Fig S2.

Because CNV could be due to differences in rDNA clusters on sex chromosomes, and our initial taxon sampling was strongly biased towards males, we included in the final dataset rDNA profiles for at least one female of each species that showed rDNA regions with inflated CN (*i.e.,* greater than a 3-fold increase in copy number of any rDNA region) to determine whether profiles were stable in both sexes. We also tested the stability of rDNA profiles across varying numbers of input reads by generating profiles for the same four specimens using 10M, 5M, and 1M reads as input into read mapping.

We tested whether rDNA profiles could be conveniently obtained as part of a hybrid capture sequencing project by simply spiking an aliquot of unenriched library into a sequencing run that included enriched libraries for the same sample. We conducted solution-based hybrid enrichment of nine breve group specimens (Table S2) using a MYcroarray (Ann Arbor, MI) custom bait set designed to target approximately 1200 loci from carabid beetles. We enriched 500 ng of *B. oromaia* library LIB0308 in an individual capture reaction. For the remaining 8 libraries, we pooled 63 ng of each library and conducted enrichment in a multiplexed capture reaction. We incubated capture reactions with biotinylated RNA baits for 16–20 hours at 65° and captured the target-hybridized baits using streptavidin-coated beads. Following manufacturer recommended wash steps, we released the enriched bead-bound targets from RNA baits via heat denaturation. We amplified enriched targets for 14 cycles using a Kapa Library Amplification Kit (Kapa Biosystems), and purified amplified products using Aline PCRClean DX beads (Aline Biosciences).

Prior to sequencing enriched libraries, we made a pool of unenriched libraries for each sample, and spiked that pool into the same lane that was used to sequence the enriched libraries such that approximately half of the reads obtained for a given sample would be from the enriched library, and half would be from the unenriched library. Because the enriched library was generated from an aliquot of the unenriched library, and therefore had the same combination of dual indices, the reads derived from both sources would be indistinguishable in the sequencing output. Importantly, the bait set used to enrich libraries lacked baits targeting rDNA regions, such that any reads contributing to rDNA profiles should have been unaffected by enrichment. This design allowed us to test the assumption that reads obtained from enriched+unenriched pooled libraries would produce effectively equivalent rDNA profiles as those produced from a dedicated sequencing run of unenriched whole genomic libraries.

Enriched libraries and their unenriched counterparts were pooled and sequenced on the same 150 base PE Illumina HiSeq3000 lane at the Oregon State Center for Genome Research and Biocomputing. We processed resulting reads in the same way as samples not subject to enrichment, except that they were down sampled to 20M reads instead of 10M reads, to account for the fact that approximately half the reads were expected to have originated from molecules belonging to the enriched libraries.

**Cytogenetic mapping of rDNA**

We performed fluorescence *in situ* hybridization (FISH) experiments with three *breve* group species, *Bembidion lividulum, B. vulcanix,* and *B. testatum*. We designed fluorescent probes to target two ~1100 base regions of the rDNA cistron in 18S and 28S respectively. The 18S target lacks copy number inflation in *B. lividulum* and *B. vulcanix*, and has minor (approximately two-fold) inflation in *B. testatum*. The 28S target region shows marked copy number inflation in *B. lividulum*, but lacks inflation in *B. vulcanix* and *B. testatum*. We hybridized probes to chromosome squashes of testis tissue taken from each species. We prepared chromosome squashes such that two squashes were available from each individual (one from each testis). This design allowed us to compare the pattern of FISH signals for both 18S and 28S in the same individual. With few exceptions, both 18S and 28S hybridizations were conducted for the same individual, in the same FISH experiment (i.e., the same batch of chemicals, incubation duration, and wash conditions). We confirmed results using multiple probe synthesis and post-hybridization wash strategies, and multiple fluorophores.

*Chromosome preparation*

We fixed tissue for chromosome squashes following Larracuente and Ferree (Larracuente and Ferree 2015). We dissected testes from freshly collected specimens of *Bembidion lividulum* and *B. testatum* in 1X PBT, and transferred tissue to 0.5% sodium citrate for 10 minutes to promote chromosome spreading. We incubated tissue in 2.5% paraformaldehyde in 45% acetic acid for four minutes on a Sigmacote (Sigma-Aldrich) treated coverslip, and squashed chromosomes onto a polylysine-coated slide by folding the slide and coverslip in filter paper and applying firm downward pressure to the coverslip for 30 seconds.

*Probe synthesis and fluorescence* in-situ *hybridization (FISH)*

We used as input for probe synthesis ~1100 base fragments of 18S and 28S (Table S4) generated via PCR. We cleaned and concentrated PCR products using Aline PCRClean DX beads (Aline Biosciences), and quantified total DNA using Qubit

Fluorometer (Life Technologies) with a Quant-iT dsDNA HS Assay Kit. We generated biotin-labeled probes using Biotin-Nick Translation Mix (Roche) following the manufacturer's recommended protocol. We verified that input DNA was fragmented between 200-700 bases using gel electrophoresis before completing probe synthesis. We purified labeled probes with Centri-Sep spin columns (Princeton Separations).

We pretreated fixed chromosome squashes with RNase and pepsin prior to FISH following Symonová *et al*., (Symonová et al. 2015). We incubated slides with 200 µl of RNase solution (200 µg of RNase in 1 ml 2X SSC buffer) for three hours at 37° and then for 3 minutes in pepisinization solution (0.005% pepsin in 0.01 N HCL) at 37°. After dehydrating slides in ethanol and air drying, we incubated slides for five minutes at 95°with 20 µl of a hybridization solution containing 100 ng of fluorescently labeled probe, 10 µl formamide, 4 µl 50 % dextran sulfate, and 2 µl 20 SSC, and 4 µl H2O. Slides were then cooled slightly, wrapped in parafilm, placed in a humidity chamber, and incubated for 14-20 hours at 37°.

In early FISH experiments we compared two post-hybridization washing approaches, a simplified protocol (Larracuente and Ferree 2015) in which slides were washed 3X for 15 min in 0.1X SSC at room temperature, and a more stringent approach in which slides were washed 3X for 5 minutes in 4X SSCT at 42°, and then 3X in 0.1X SSC at 60° for 5 minutes (Larracuente 2017). Although consistent patterns of FISH signals were observed regardless of wash strategy, we used the latter, more stringent protocol (Larracuente 2017) for results reported herein as it resulted in a better signal to noise ratio. Following post-hybridization washes, we incubated slides with 100 µl blocking solution for 30 minutes at 37°. Following blocking we detected probes by incubation with Streptavidin, Rhodamine Red™-X conjugate (ThermoFisher) for 30 minutes at 37°, and repeated the post-hybridization wash steps as described above before mounting slides following Larracuente (Larracuente 2017).

We counterstained and mounted FISH slides using 11 µl of SlowFade® Diamond Antifade Mountant with DAPI (ThermoFisher) and imaged FISH signals using a Zeiss LSM 780 NLO Confocal Microscope System. We verified FISH patterns resulting from the above protocol on dozens of nuclei from five *B. lividulum* and *B. vulcanix* individuals, and three *B. testatum* individuals. We also verified *B. lividulum*-specific FISH patterns on condensed chromosomes in at least 10 nuclei from five replicate individuals. We tested for non-specific binding of 28S probes generated from *B. testatum* and *B. testatum* PCR products by hybridizing probes generated from *B. lividulum* PCR products to *B. testatum* chromosomes, and hybridizing *B. testatum*-generated probes to chromosomes of *B. lividulum*.

In addition to our preferred protocol described above we tested for the presence of probe-specific, and fluorophore-specific artifacts in FISH patterns by using alternative probe synthesis and FISH approaches. We designed primers to amplify two non-overlapping ~500 base amplicons in each target gene (Table S4 Figs. 7 and S25). We pooled PCR products by locus such that both 18S amplicons were in a pool and both 28S products were in pool. We cleaned and quantified PCR products as described above, and directly labeled 1µg of DNA from each pool using a ULYSIS® Alexa Fluor® 488 Nucleic Acid Labeling Kit (ThermoFisher) following the manufacturer's recommended protocol. Sample preparation and hybridization of these probes followed our preferred protocol described above, except that it eliminated blocking and detection following the post-hybridization washes and the probes were directly labeled with fluorophores.

**Testing for rDNA profile variation across Bembidion (Plataphus)**

We tested for profile variation across a broader taxonomic scope by sampling 41 species across the subgenus *Plataphus* (in addition to the nine *breve* group species) (Table S5). We selected multiple taxa from each major clade in the subgenus, including all known species within several groups of closely related species.

We inferred the *Plataphus* phylogeny in IQ-TREE v6.5 (Nguyen et al. 2014) as orchestrated by Mesquite v3.5 (Maddison WP and Maddison DR 2018) and Zephyr v2.1 (Maddison DR and Maddison WP 2018) using a six-gene dataset consisting of previously published data (Maddison 2008, 2012; Hildebrandt and Maddison 2011; Maddison and Ober 2011; Sproul and Maddison 2017b, 2017a; Maddison and Maruyama 2018) and sequences new to this study (Table S6). The latter have been submitted to GenBank with accession numbers MK461576 – MK461828.

We assembled chromatograms using Phred (Green and Ewing 2002) and Phrap (Green 1999) via the Chromaseq package in Mesquite v3.2 (Maddison and Maddison 2014, 2015). Final sequence editing was conducted manually in Chromaseq. We aligned sequences from protein-coding genes in Mesquite; no insertion or deletion events need be presumed in the history of the sequences examined. The ribosomal gene (28S) was aligned in MAFFT 7.130b (Katoh and Toh 2008) with the G-INS-I algorithm as implemented in Mesquite. Following alignment, data matrices for each gene were prepared for downstream analysis using Mesquite. We generated rDNA profiles for *Plataphus* species using the same methods described above, except that we relaxed the stringency of read mapping parameters slightly (length fraction=0.80 and similarity fraction=0.80) following the results of our parameter sensitivity analysis (Fig. S2).

We used IQ-TREE to find optimal models of character evolution and partitioning of the data, and for tree inference. The beginning partition had each codon position in each gene as a separate partition, plus all of 28S in another partition (Table S7). We searched for the Maximum Likelihood tree across 100 search replicates.

**Cluster analysis of repetitive DNA**

We further validated patterns observed in rDNA profiles, and characterized variation of other DNA repeats using RepeatExplorer (Novák et al. 2010). RepeatExplorer uses short-read sequence data to generate graph-based clusters (Blondel et al. 2008) of assembled repeats, and annotates clusters using public databases (Jurka et al. 2005; Marchler-Bauer et al. 2010). We conducted cluster analysis on all but eight *breve* group specimens for which rDNA profiles were generated as part of a hybrid capture workflow (Table S2), as the inclusion of enriched loci violates assumptions of the analysis. Prior to cluster analysis we estimated genomic coverage and down sampled reads to 0.25x coverage in CLC GW, and conducted clustering using the RepeatExplorer Galaxy-based web server (Novák et al. 2013). RepeatExplorer output orders clusters based on genome proportion. We analyzed the top 100 clusters (i.e., the 100 clusters most abundant in the genome) and grouped clusters into six repetitive DNA categories: Class I transposable elements (TEs), Class II TEs, rDNA, simple repeats, unknown clusters, and unknown clusters containing rDNA hits from BLAST and RepeatMasker databases. We generated pie charts to visualize variation in repetitive DNA across specimens. For *Bembidion lividulum* we also analyzed within-species variation in Class I and Class II TE abundance. We scanned cluster analysis results for TE protein domains using the Protein Domain Finder tool in RepeatExplorer and plotted the total number of clusters containing hits for each of nine TE superfamilies for all *B. lividulum* individuals.

**Generating rDNA profiles using freely available software tools**

As an alternative approach to generating rDNA profiles in CLC GW, we indexed the *Bembidion aeruginosum* rDNA reference sequence and conducted read mapping of Illumina reads in Bowtie2 v3.2.4.2 (Langmead and Salzberg 2012) using the 'bowtie2-build' command. We used SAMtools v1.9 (Li et al. 2009) to convert read mapping output to BAM format, sort and index the resulting BAM files using the 'sort' and 'index' functions, and generate a table of read depth at each position using the 'depth' function with flags '–a' to retain 0-value positions, and '–d=0' to avoid capping coverage values at 8000. We generated rDNA profiles in R v3.5.1 (R Core Team 2013) by reading in the table of read depth values using the 'read.table' command and making a barplot of coverage values using the 'barplot' function, which produces a plot that can be saved as a vector file.

### DATA ACCESSIBILITY

Sequence read files are deposited in NCBI Sequence Read Archive (Accessions SAMN10860544– SAMN10860633). PCR-based sequence data are deposited in GenBank (Accessions MK461576 – MK461828). Matrices and results from phylogenetic analysis, as well as the reference sequence file used for generating rDNA profiles are deposited in DRYAD (entry doi: XXX) (to be added upon acceptance).

### SUPPLEMENTAL FIGURE CAPTIONS

**Figure S1.** Images of nine species in *breve* species group of *Bembidion* (Carabidae). (A) *Bembidion lividulum*; (B) *B. breve*; (C) *B. testatum*; (D) *B. saturatum*; (E) *B. vulcanix*; (F) *B. geopearlis*; (G) *B. oromaia*; (H) *B. laxatum*; (I) *B. ampliatum*.

**Figure S2.** Parameter sensitivity analysis. An example of profiles generated for the same specimen (*Bembidion sp.nr.curtulatum "Idaho"* 2145) using read mapping parameters of varying stringency with A being the most stringent and F being the least stringent. Parameter settings used are described in the text and Table S3. The settings selected for final analysis are indicated by the arrow.

**Figure S3.** Reference sequence sensitivity analysis. An example of rDNA profiles illustrating the difference between a profile generated by mapping reads to a reference sequence derived from a closely related taxon compared to a profile generated by mapping reads to a reference sequence derived from a conspecific sample. Red arrows indicate peaks due to CN variation that are present regardless of reference choice. Black arrows indicate valleys that are artifacts that can be present when mapping reads to a non-conspecific reference, and presumably arise due to sequence divergence (including indels) between the reference sequence and the reads being mapped. The blue arrow indicates a small peak, an artifact of read mapping, that can be appear adjacent to a large valley. The position of rRNA genes relative to rDNA profiles is shown along the bottom for reference.

**Figure S4.** rDNA profiles from nine *breve* group species as follows: (A) *Bembidion vulcanix* (4649); (B) *B. oromaia* (4250); (C) *B. laxatum* (5086); (D) *B. ampliatum* (4245); (E) *B. saturatum* (3313); (F) *B. geopearlis* (4731); (G) *B. testatum* (4169); (H) *B. breve* (4187); (I) *B. lividulum* (3486). The position of rRNA genes relative to rDNA profiles is shown along the bottom for reference.

**Figure S5.** The *Bembidion lividulum* clade taken from the species tree in Fig. 3 with rDNA profiles. Features in rDNA profiles that appear to show phylogenetic signal are indicated with arrows and text. Profiles generated from female specimens are indicated by the female symbol. Pie charts indicate the fraction of clusters in the top 100 clusters that belong to each of six categories of repetitive DNA: Class I TEs in blue, Class II TEs in red, simple repeats in green, rDNA in orange, and unknown with rDNA hits in pink. Repeat categories are also summarized in a visual legend that applies to Figs. S5–S13.

**Figure S6.** The *Bembidion saturatum* clade taken from the species tree in Fig. 3 with rDNA profiles. Profiles generated from female specimens are indicated by the female symbol. Two morphologically distinct specimens suspected of belonging to cryptic lineages in Sproul and Maddison (2017a) are indicated by a red star. Pie charts show the fraction of repetitive DNA in each of six categories explained further in the Fig. S5 caption.

**Figure S7.** The *Bembidion ampliatum* clade taken from the species tree in Fig. 3 with rDNA profiles. Profiles generated from female specimens are indicated by the female symbol, and those generated from hybrid enrichment sequencing are annotated with

“HybSeq”. Pie charts show the fraction of repetitive DNA in each of six categories explained further in the Fig. S5 caption.

**Figure S8.** The *Bembidion breve* clade taken from the species tree in Fig. 3 with rDNA profiles. Profiles generated from female specimens are indicated by the female symbol. Pie charts show the fraction of repetitive DNA in each of six categories explained further in the Fig. S5 caption.

**Figure S9.** The *Bembidion geopearlis* clade taken from the species tree in Fig. 3 with rDNA profiles. Profiles generated from female specimens are indicated by the female symbol, and those generated from hybrid enrichment sequencing are annotated with “HybSeq”. Pie charts show the fraction of repetitive DNA in each of six categories explained further in the Fig. S5 caption.

**Figure S10.** The *Bembidion laxatum* clade taken from the species tree in Fig. 3 with rDNA profiles. Profiles generated from female specimens are indicated by the female symbol. Pie charts show the fraction of repetitive DNA in each of six categories explained further in the Fig. S5 caption.

**Figure S11.** The *Bembidion oromaia* clade taken from the species tree in Fig. 3 with rDNA profiles. Profiles generated hybrid enrichment sequencing are annotated with “HybSeq”. Pie charts show the fraction of repetitive DNA in each of six categories explained further in the Fig. S5 caption.

**Figure S12.** The *Bembidion testatum* clade taken from the species tree in Fig. 3 with rDNA profiles. Profiles generated from female specimens are indicated by the female symbol, and those generated from hybrid enrichment sequencing are annotated with “HybSeq”. Pie charts show the fraction of repetitive DNA in each of six categories explained further in the Fig. S5 caption.

**Figure S13.** The *Bembidion vulcanix* clade taken from the species tree in Figure? with rDNA profiles. Profiles generated hybrid enrichment sequencing are annotated with “HybSeq”. Pie charts show the fraction of repetitive DNA in each of six categories explained further in the Fig. S5 caption.

**Figure S14.** Species distribution map for *Bembidion lividulum* showing the geographic sampling of rDNA profiles. Localities from with specimens belonging to Clade 1 are shown by circles, while localities with specimens belonging to Clade 1 are shown by stars. Circles and stars outlined in red indicate localities from which we obtained rDNA profiles, with numbers in shapes corresponding to rDNA profiles shown on the left of the figure. Features in rDNA profiles that appear to show geographic signal are indicated with arrows and text.

**Figure S15.** Species distribution map for *Bembidion saturatum* showing the geographic sampling of rDNA profiles. Confirmed localities of the species are shown by either small black circles, larger red-outlined black circles, or black stars with a red outline. Red-outlined circles and stars indicate those localities from which we obtained rDNA profiles. Localities containing morphologically distinct specimens suspected of belonging to cryptic lineages in Sproul and Maddison (2017a) are indicated red-outlined stars. Numbers in circles and stars correspond to rDNA profiles shown on the left of the figure.

**Figure S16.** Species distribution map for *Bembidion ampliatum* showing the geographic sampling of rDNA profiles. Confirmed localities of the species are shown by either small black circles, or larger red-outlined black circles, the latter indicating those localities from which we obtained rDNA profiles. Numbers in red-outlined circles correspond to rDNA profiles shown on the left of the figure.

**Figure S17.** Species distribution map for *Bembidion breve* showing the geographic sampling of rDNA profiles. See the figure caption for Fig. S16 or additional explanation.

**Figure S18.** Species distribution map for *Bembidion geopearlis* showing the geographic sampling of rDNA profiles. See the figure caption for Fig. S16 for additional explanation.

**Figure S19.** Species distribution map for *Bembidion laxatum* showing the geographic sampling of rDNA profiles. See the figure caption for Fig. S16 for additional explanation.

**Figure S20.** Species distribution map for *Bembidion oromaia* showing the geographic sampling of rDNA profiles. See the figure caption for Fig. S16 for additional explanation.

**Figure S21.** Species distribution map for *Bembidion testatum* showing the geographic sampling of rDNA profiles. See the figure caption for Fig. S16 for additional explanation.

**Figure S22.** Species distribution map for *Bembidion vulcanix* showing the geographic sampling of rDNA profiles. See the figure caption for Fig. S16 for additional explanation.

**Figure S23.** A histogram of average rho values within a given species resulting from the correlation analysis between rDNA profiles and species boundaries.

**Figure S24.** rDNA profiles obtained from the same specimen (*B. lividulum* 3486) using: (A) 10 million, (B) 5 million, and (C) 1 million reads.

**Figure S25.** FISH signals obtained by cytogenetic mapping of rDNA in *B. testatum*. (A) FISH signals resulting from hybridization of 18S probes to *B. testatum* nuclei. (B) FISH signals resulting from hybridization of 28S probes to *B. testatum* nuclei. (C) a rDNA profile for *B. testatum* in standard scale below a rescaled version of the same profile shown to show the pattern of slight copy number inflation in the 18S region. The location of 18S and 28S FISH probes are indicated with green boxes.

**Figure S26.** FISH signals resulting from hybridization of 28S probes to condensed meiotic chromosomes in *B. lividulum* with arrows indicating several euchromatic tails that are free from FISH signals.

**Figure S27.** rDNA profile of 100-year-old type specimen (*B. lividulum* Casey) (G) alongside rDNA profiles of DNA-preserved specimens of *Bembidion laxatum*, (A–B); *B. ampliatum*, (C–D); and *B. lividulum* (E–F). Adapted from Sproul and Maddison (2017a).

Fig. S1

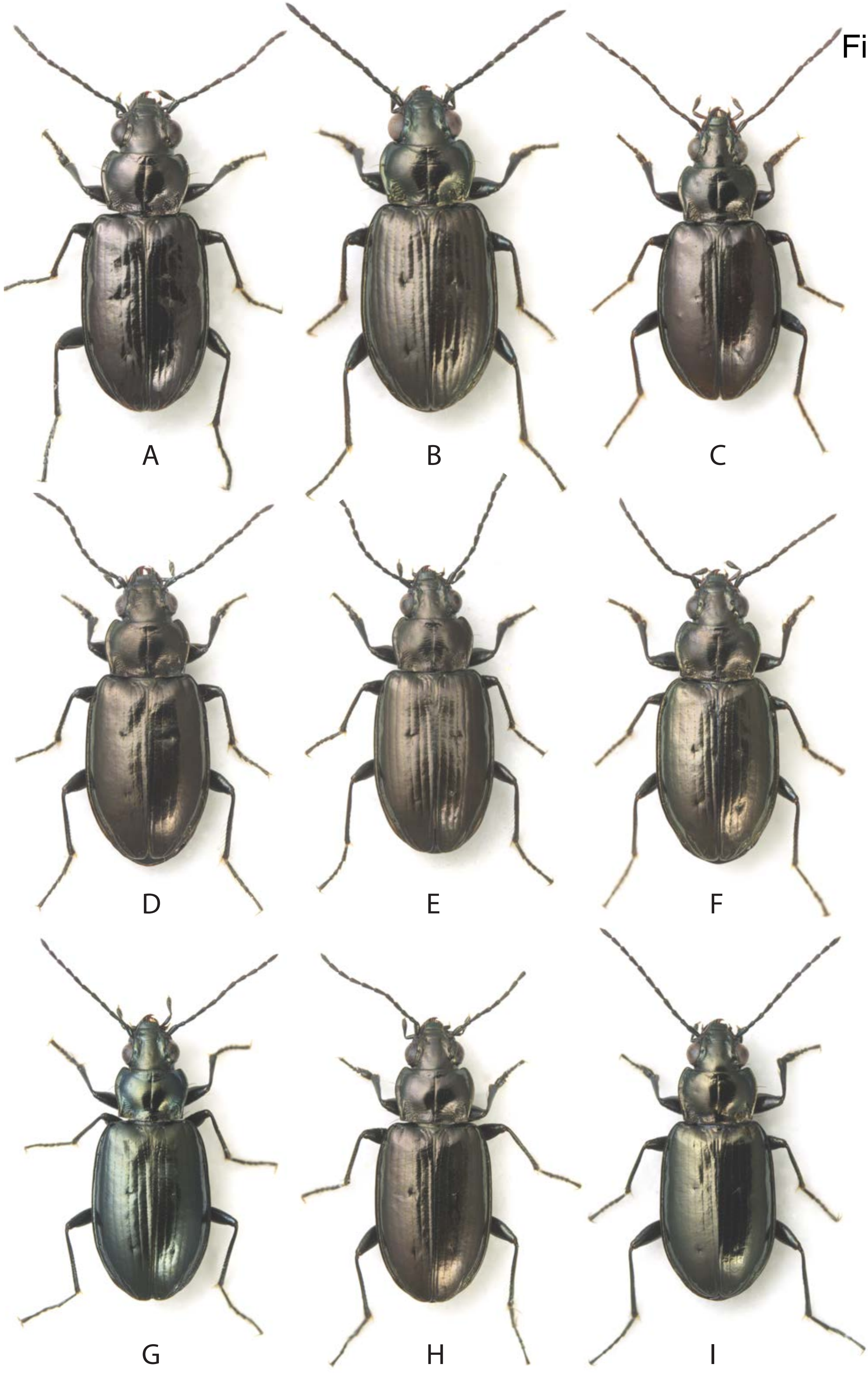

Fig. S2

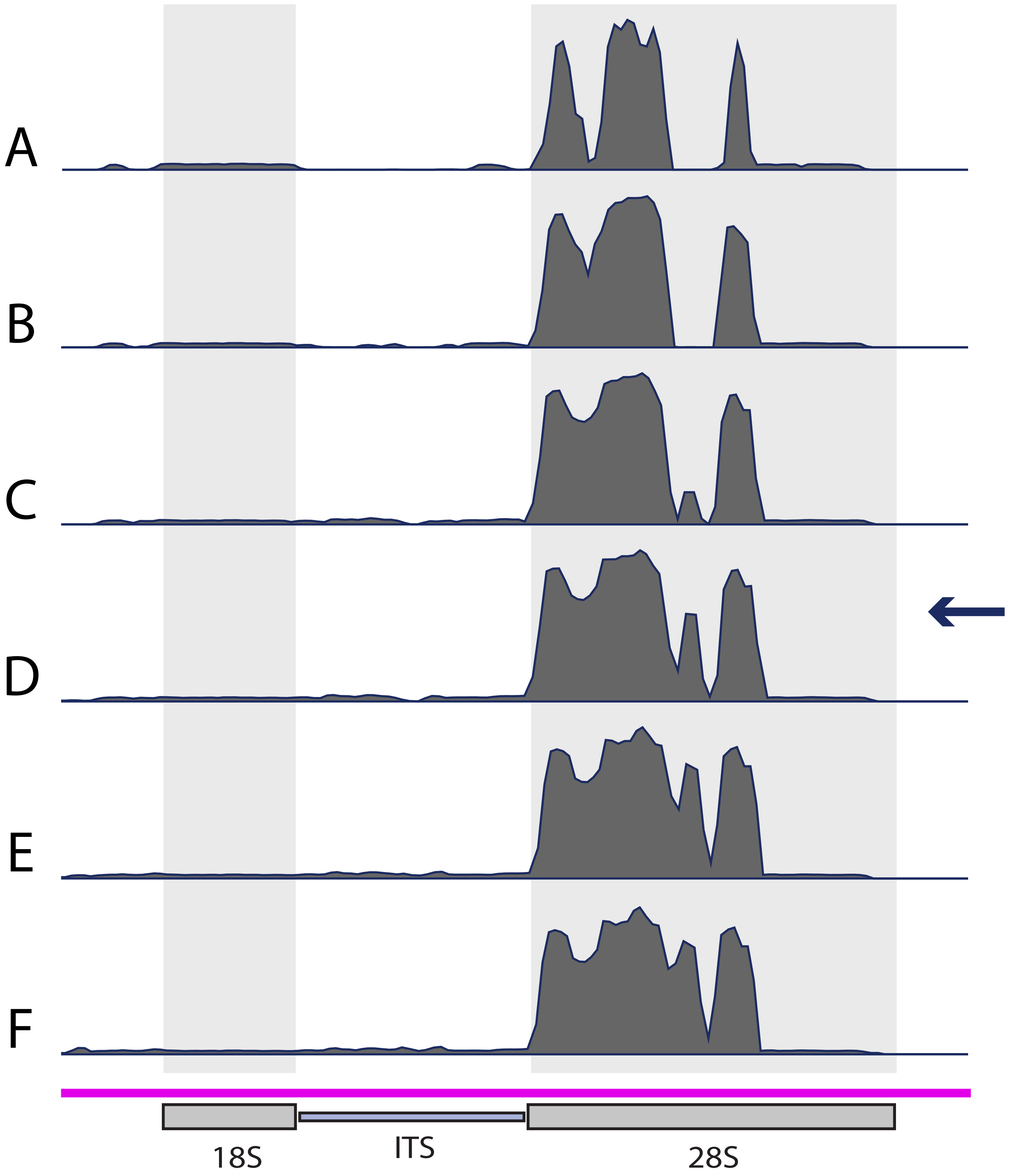

Fig. S3

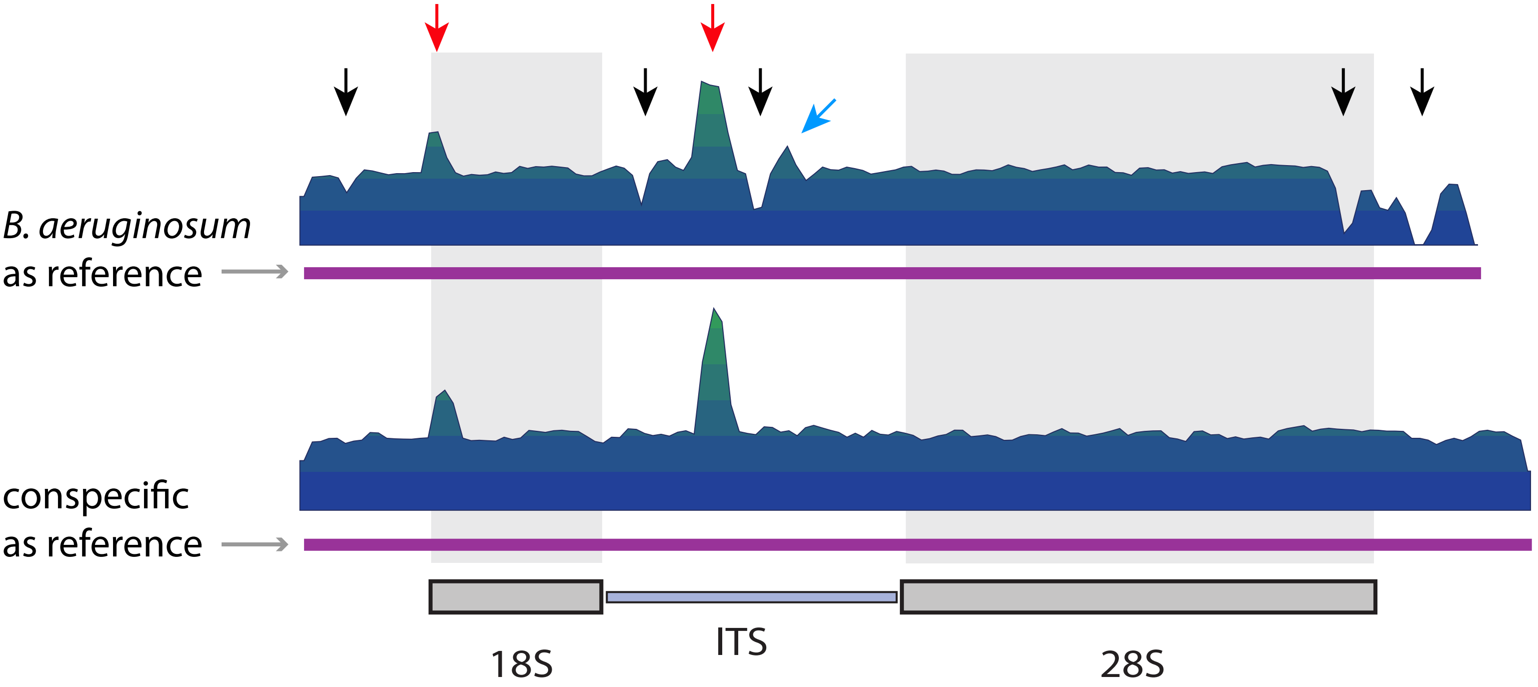
B. aeruginosum
as reference
conspecific
as reference
18S
ITS
28S

Fig. S4

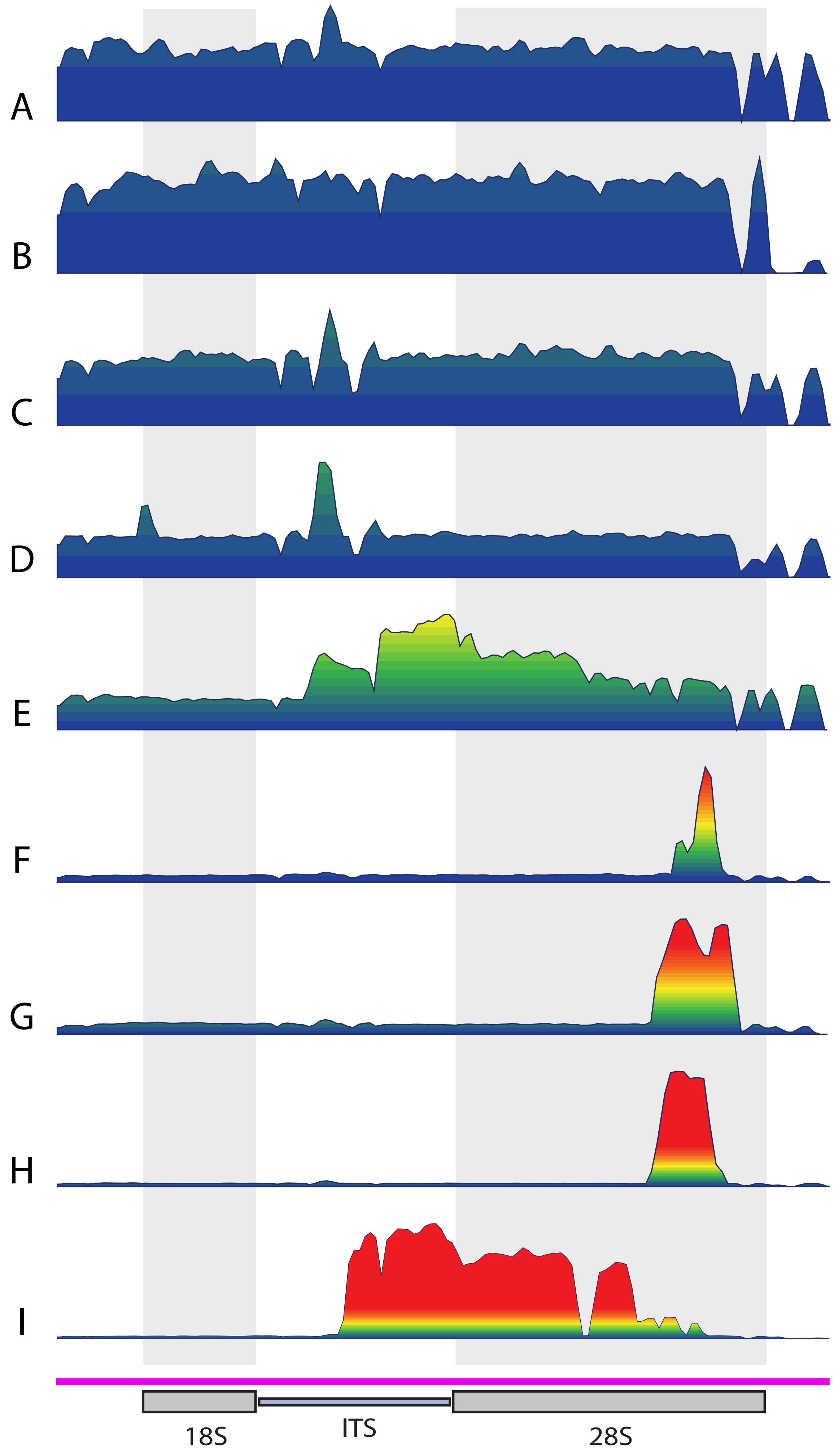

Fig. S5

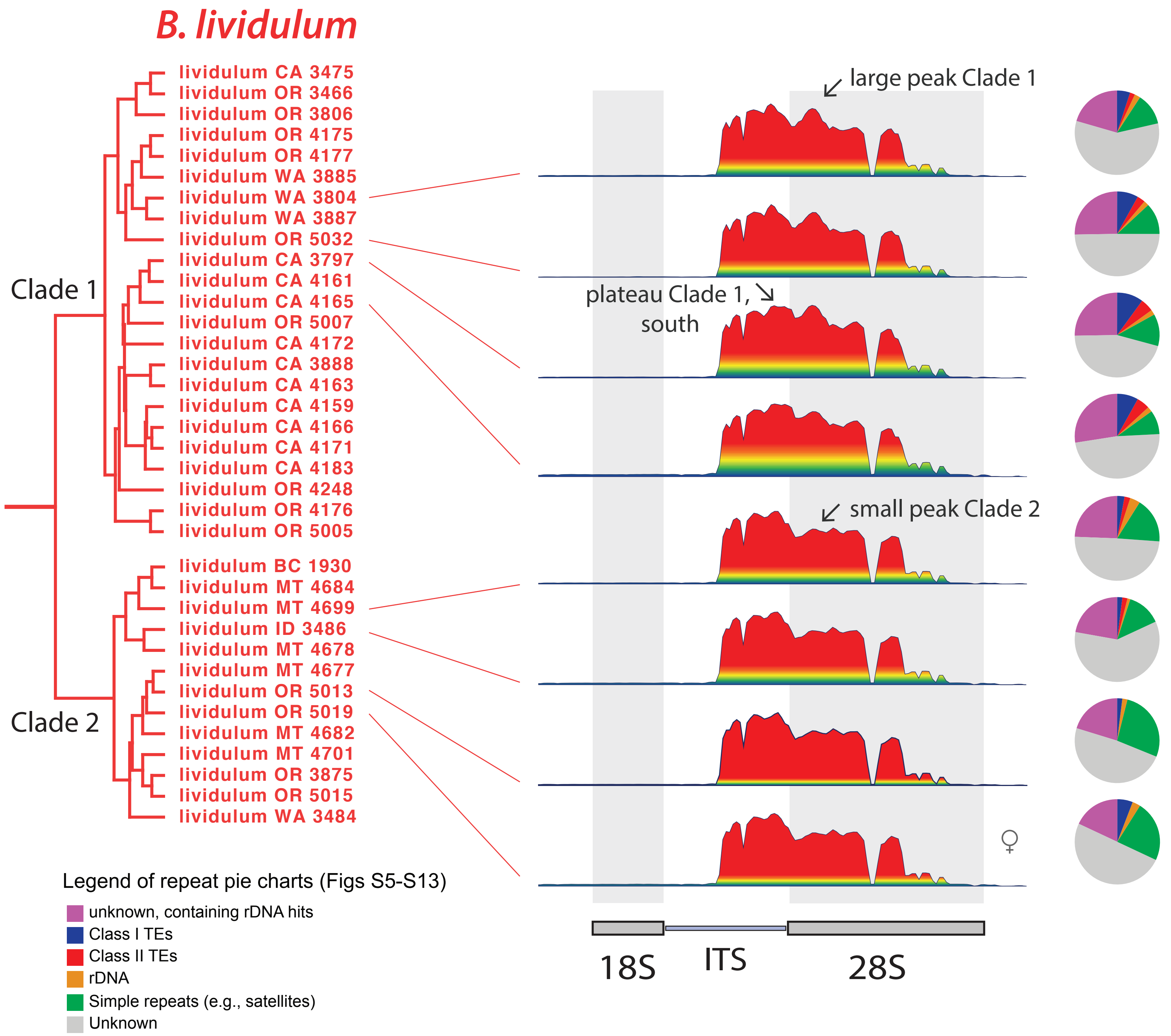
B. lividulum
lividulum CA 3475
lividulum OR 3466
lividulum OR 3806
lividulum OR 4175
lividulum OR 4177
lividulum WA 3885
lividulum WA 3804
lividulum WA 3887
lividulum OR 5032
lividulum CA 3797
lividulum CA 4161
lividulum CA 4165
lividulum OR 5007
lividulum CA 4172
lividulum CA 3888
lividulum CA 4163
lividulum CA 4159
lividulum CA 4166
lividulum CA 4171
lividulum CA 4183
lividulum OR 4248
lividulum OR 4176
lividulum OR 5005
lividulum BC 1930
lividulum MT 4684
lividulum MT 4699
lividulum ID 3486
lividulum MT 4678
lividulum MT 4677
lividulum OR 5013
lividulum OR 5019
lividulum MT 4682
lividulum MT 4701
lividulum OR 3875
lividulum OR 5015
lividulum WA 3484
Clade 1
Clade 2
large peak Clade 1
plateau Clade 1, south
small peak Clade 2
♀
18S
ITS
28S
Legend of repeat pie charts (Figs S5-S13)
unknown, containing rDNA hits
Class I TEs
Class II TEs
rDNA
Simple repeats (e.g., satellites)
Unknown

Fig. S6

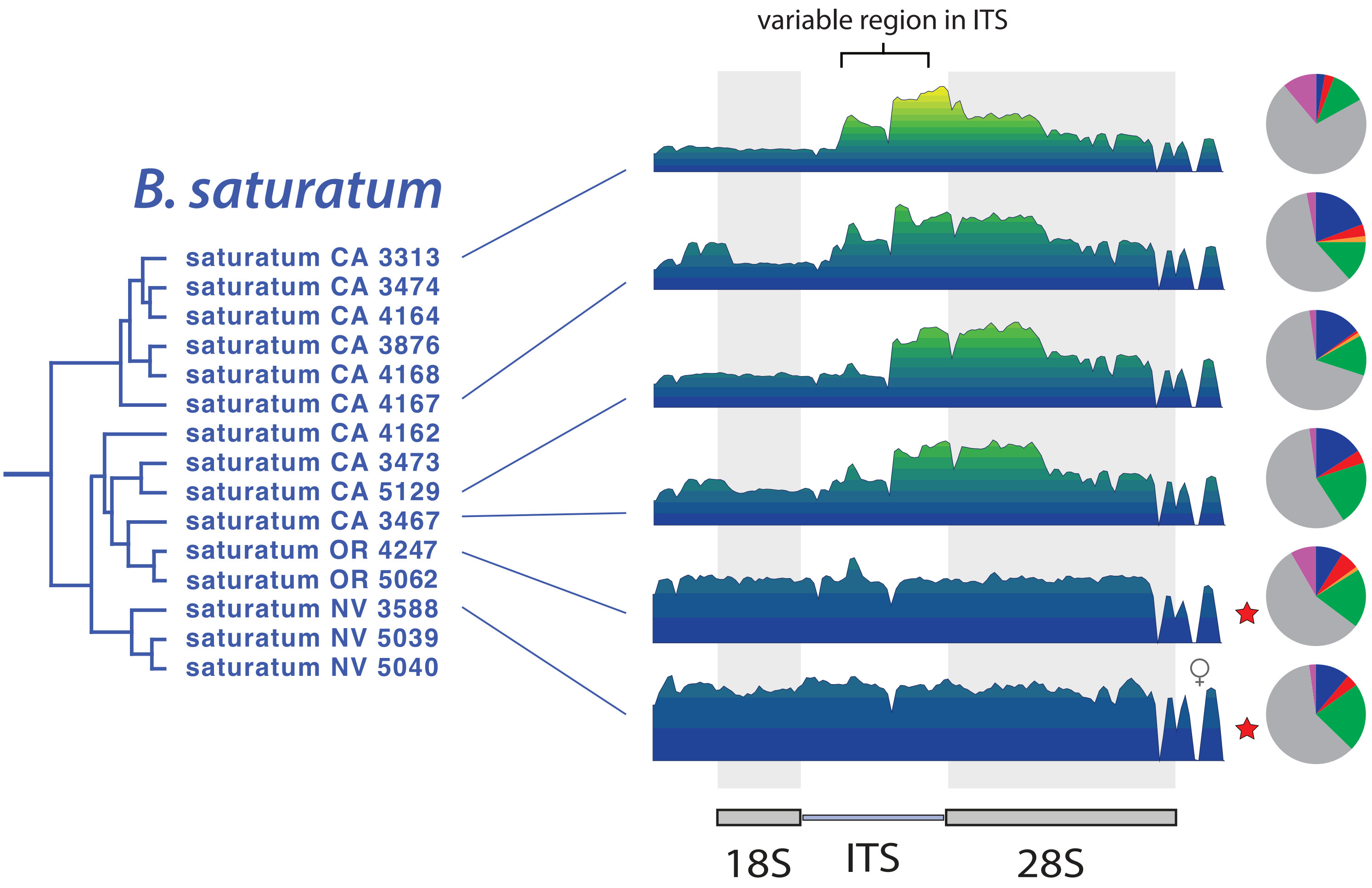

variable region in ITS
B. saturatum
saturatum CA 3313
saturatum CA 3474
saturatum CA 4164
saturatum CA 3876
saturatum CA 4168
saturatum CA 4167
saturatum CA 4162
saturatum CA 3473
saturatum CA 5129
saturatum CA 3467
saturatum OR 4247
saturatum OR 5062
saturatum NV 3588
saturatum NV 5039
saturatum NV 5040
♀
18S
ITS
28S

Fig. S7

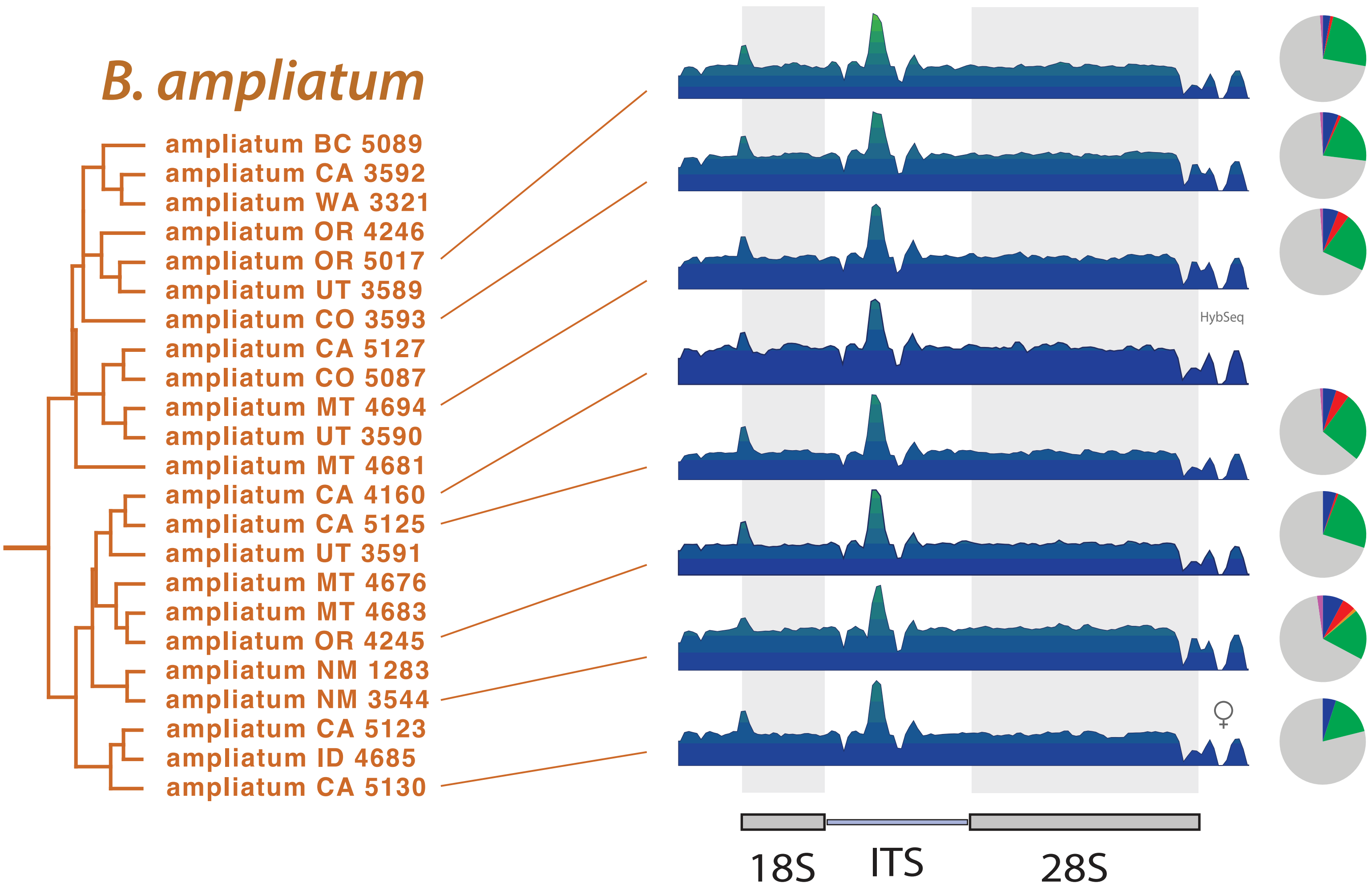

B. ampliatum
ampliatum BC 5089
ampliatum CA 3592
ampliatum WA 3321
ampliatum OR 4246
ampliatum OR 5017
ampliatum UT 3589
ampliatum CO 3593
ampliatum CA 5127
ampliatum CO 5087
ampliatum MT 4694
ampliatum UT 3590
ampliatum MT 4681
ampliatum CA 4160
ampliatum CA 5125
ampliatum UT 3591
ampliatum MT 4676
ampliatum MT 4683
ampliatum OR 4245
ampliatum NM 1283
ampliatum NM 3544
ampliatum CA 5123
ampliatum ID 4685
ampliatum CA 5130
HybSeq
♀
18S
ITS
28S

Fig. S8

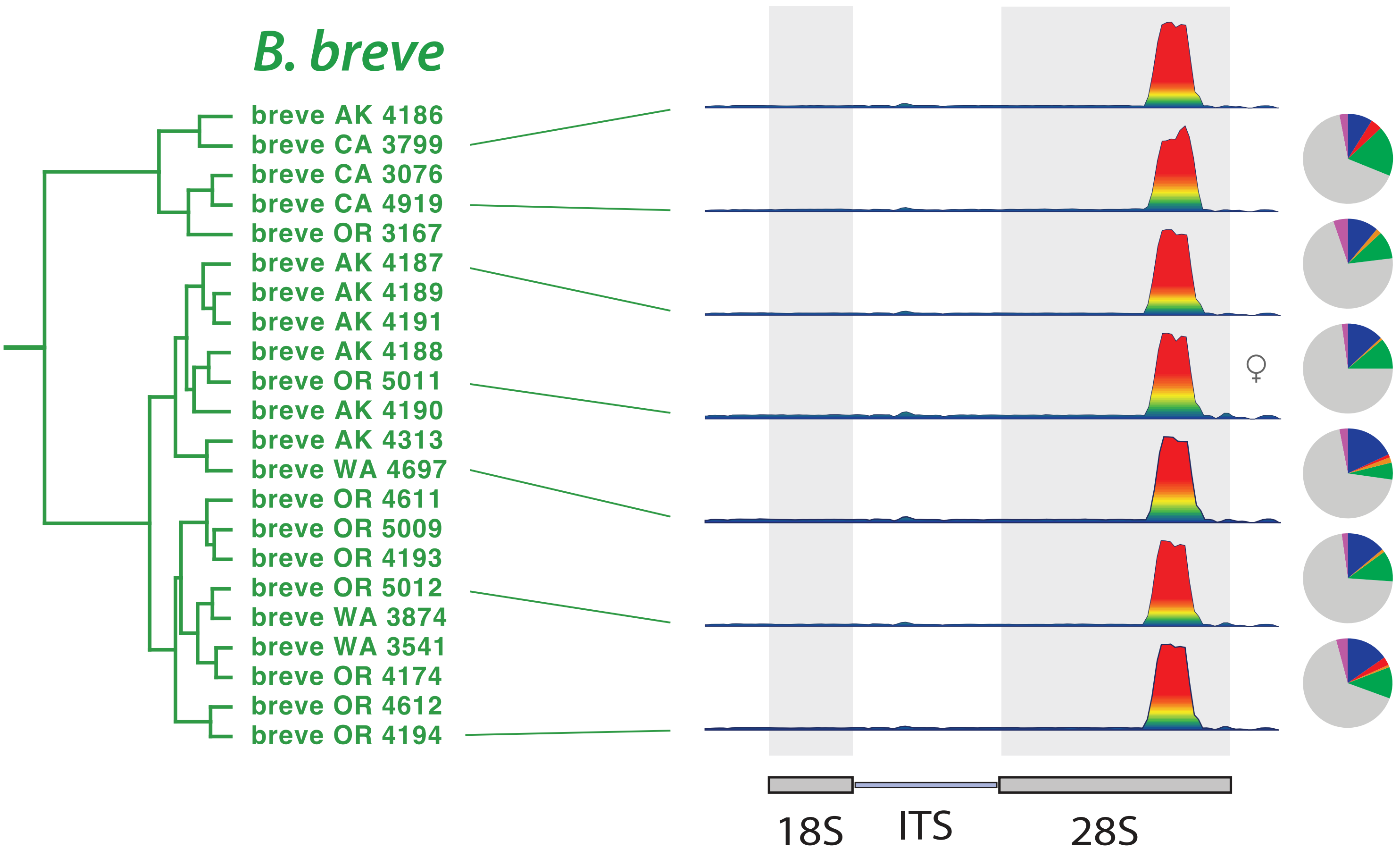
B. breve
breve AK 4186
breve CA 3799
breve CA 3076
breve CA 4919
breve OR 3167
breve AK 4187
breve AK 4189
breve AK 4191
breve AK 4188
breve OR 5011
breve AK 4190
breve AK 4313
breve WA 4697
breve OR 4611
breve OR 5009
breve OR 4193
breve OR 5012
breve WA 3874
breve WA 3541
breve OR 4174
breve OR 4612
breve OR 4194
♀
18S
ITS
28S

Fig. S9

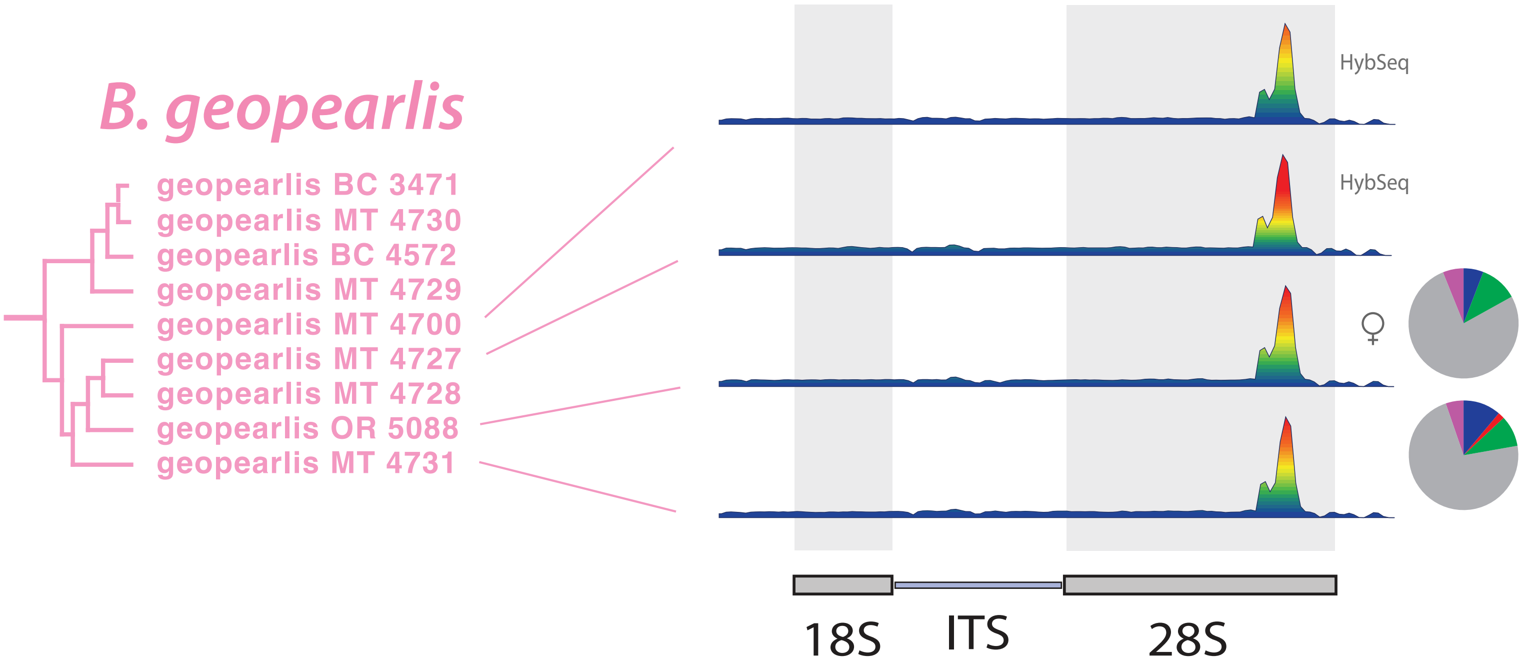

B. geopearlis
geopearlis BC 3471
geopearlis MT 4730
geopearlis BC 4572
geopearlis MT 4729
geopearlis MT 4700
geopearlis MT 4727
geopearlis MT 4728
geopearlis OR 5088
geopearlis MT 4731
HybSeq
HybSeq
♀
18S
ITS
28S

Fig. S10

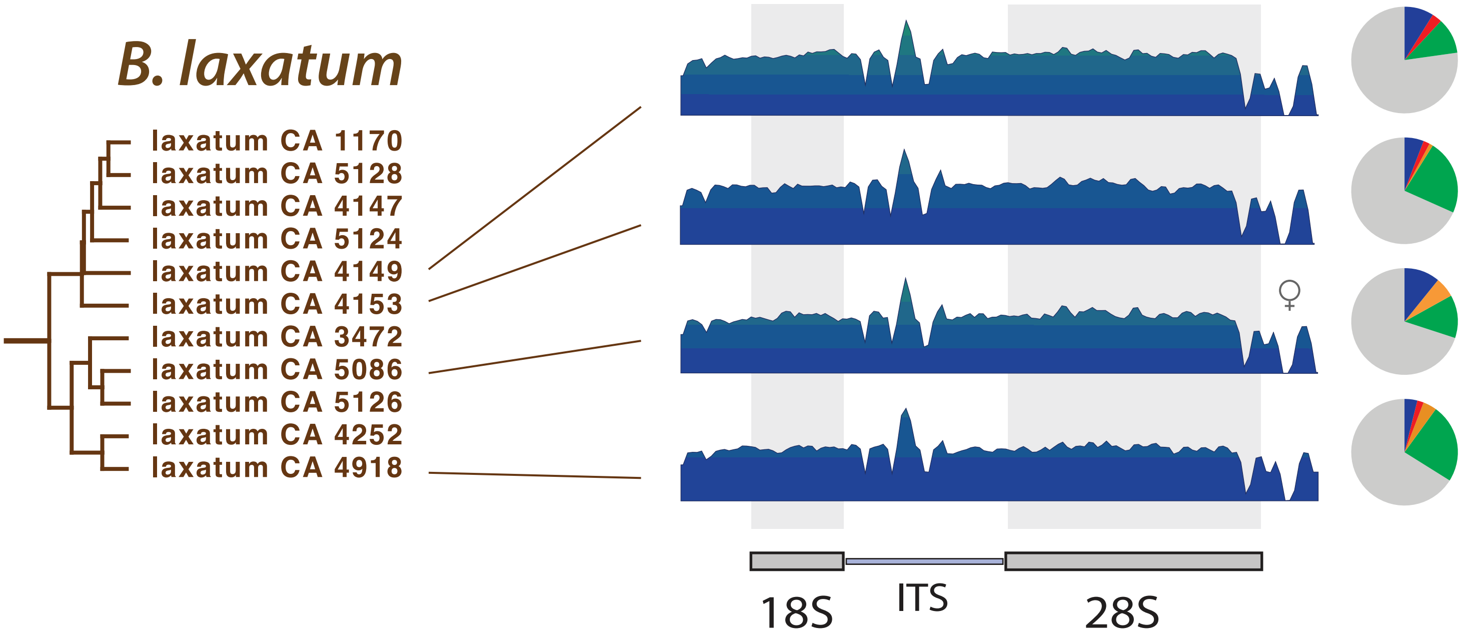
B. laxatum
laxatum CA 1170
laxatum CA 5128
laxatum CA 4147
laxatum CA 5124
laxatum CA 4149
laxatum CA 4153
laxatum CA 3472
laxatum CA 5086
laxatum CA 5126
laxatum CA 4252
laxatum CA 4918
♀
18S
ITS
28S

Fig. S11

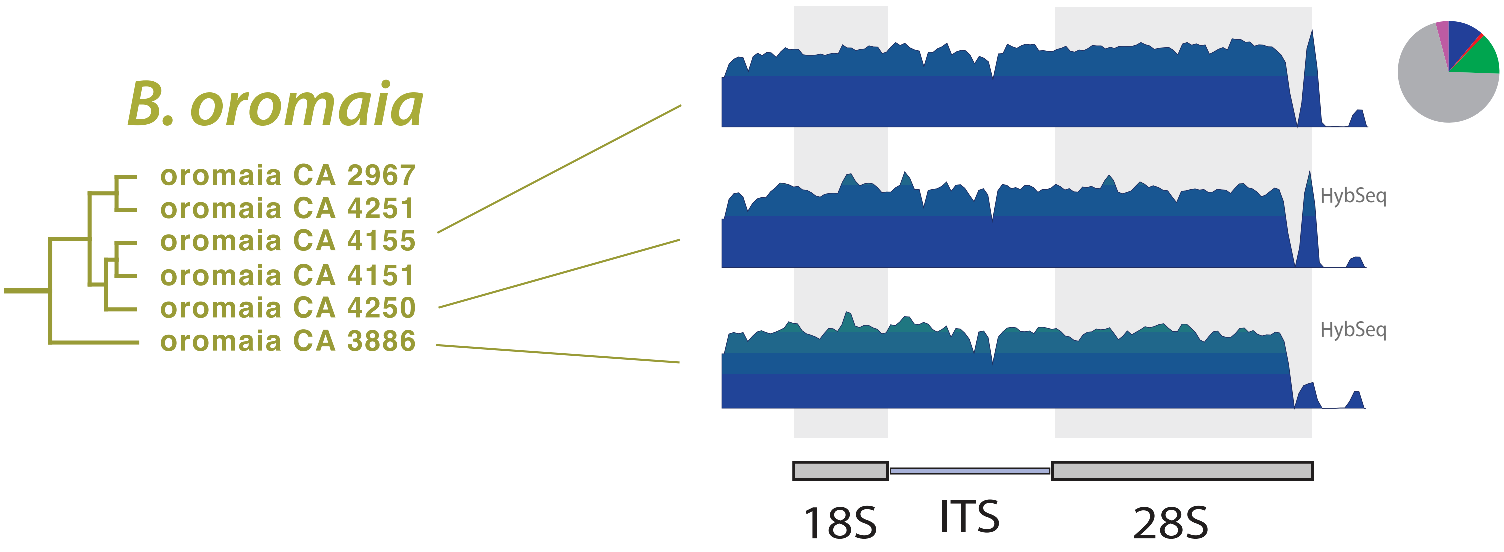
B. oromaia
oromaia CA 2967
oromaia CA 4251
oromaia CA 4155
oromaia CA 4151
oromaia CA 4250
oromaia CA 3886
HybSeq
HybSeq
18S
ITS
28S

Fig. S12

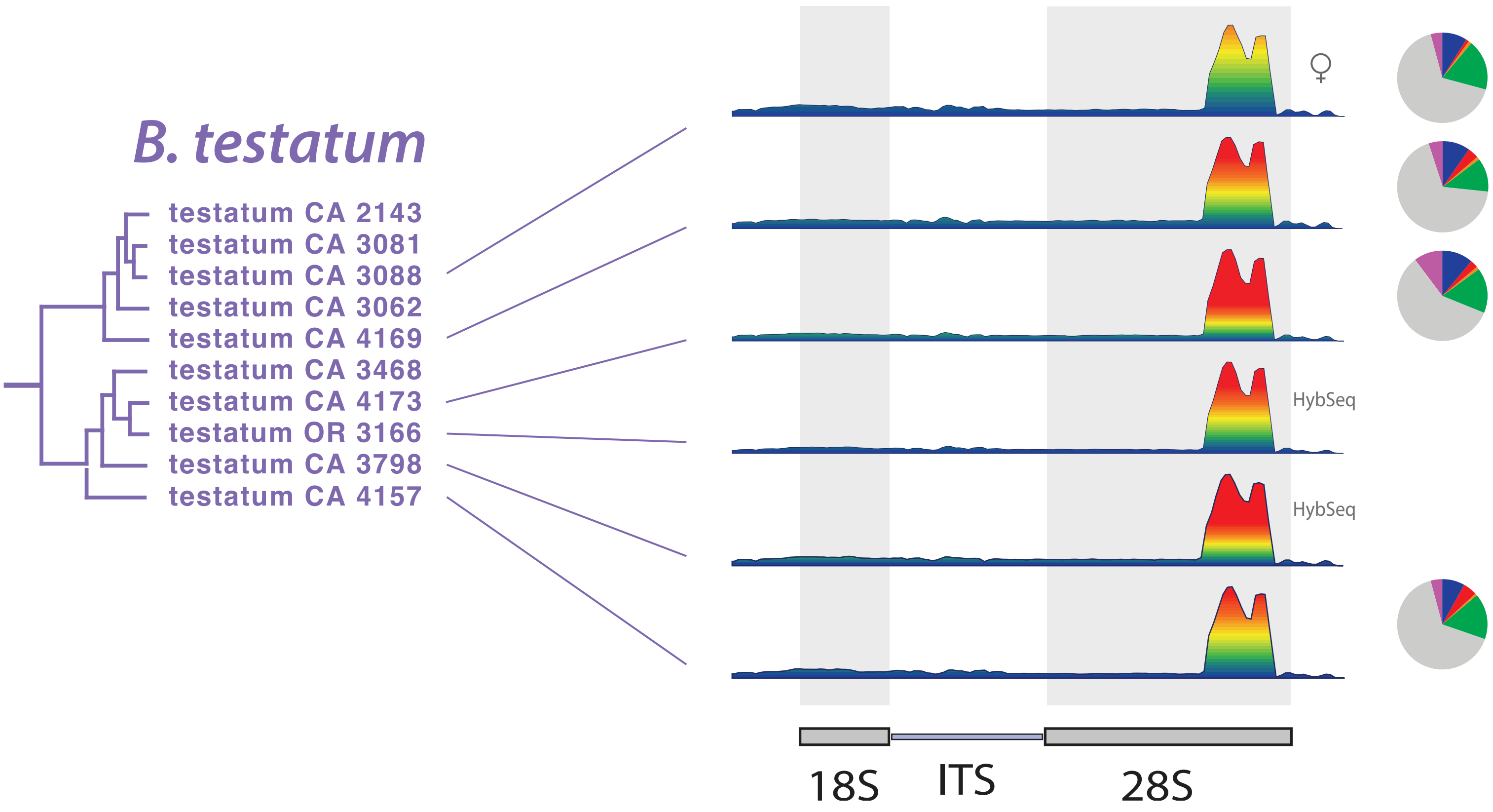
B. testatum
testatum CA 2143
testatum CA 3081
testatum CA 3088
testatum CA 3062
testatum CA 4169
testatum CA 3468
testatum CA 4173
testatum OR 3166
testatum CA 3798
testatum CA 4157
♀
HybSeq
HybSeq
18S
ITS
28S

Fig. S13

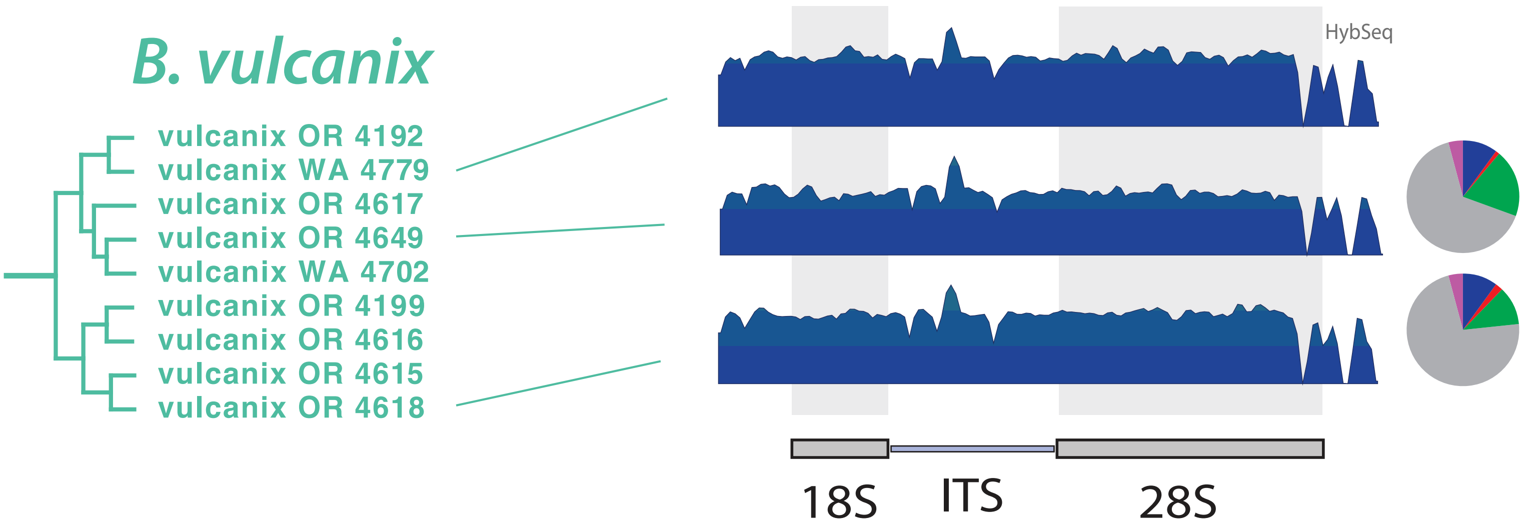

B. vulcanix
vulcanix OR 4192
vulcanix WA 4779
vulcanix OR 4617
vulcanix OR 4649
vulcanix WA 4702
vulcanix OR 4199
vulcanix OR 4616
vulcanix OR 4615
vulcanix OR 4618
HybSeq
18S
ITS
28S

Fig. S14

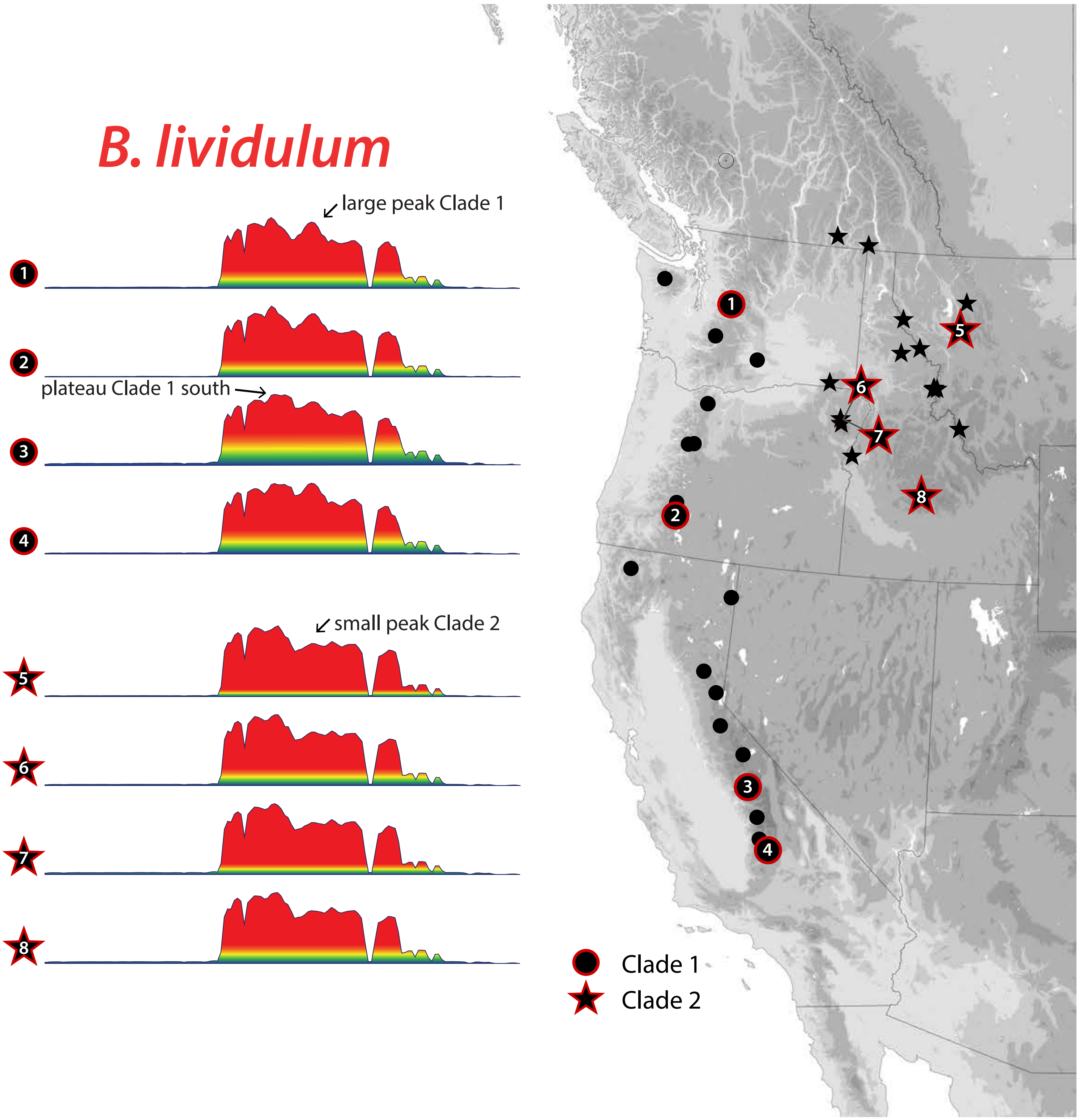
B. lividulum
large peak Clade 1
plateau Clade 1 south
small peak Clade 2
1
2
3
4
5
6
7
8
Clade 1
Clade 2

Fig. S15

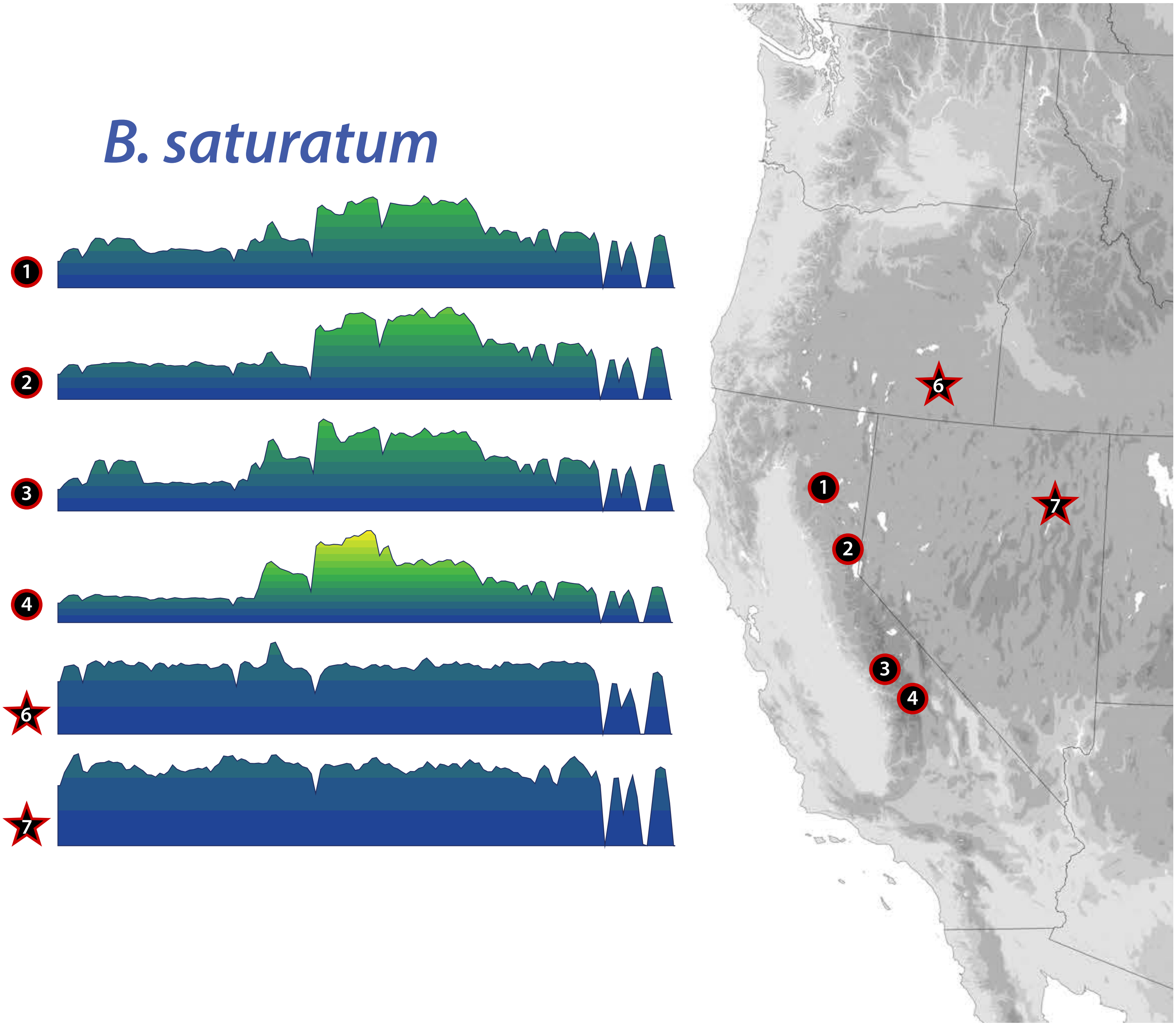

Fig. S16

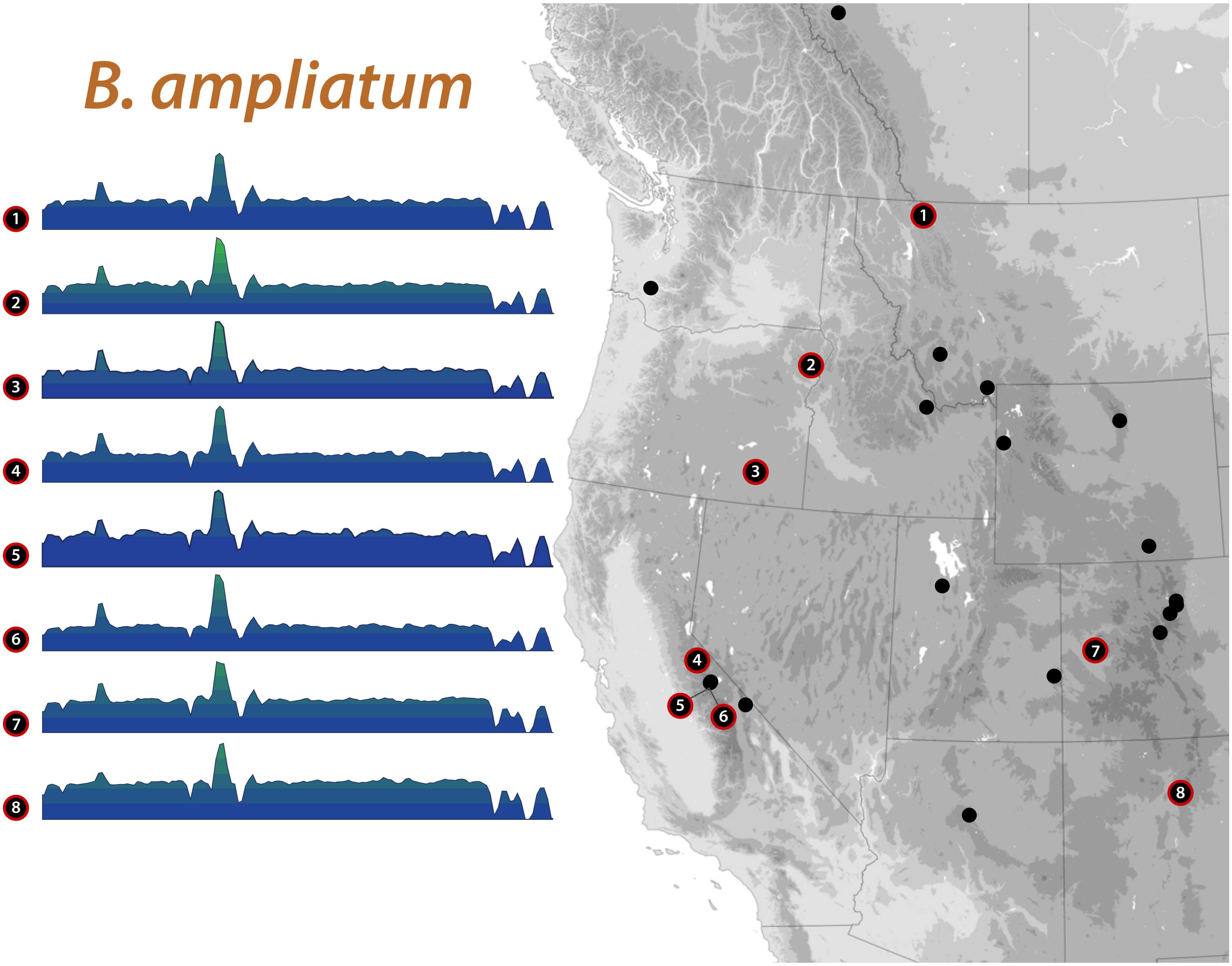

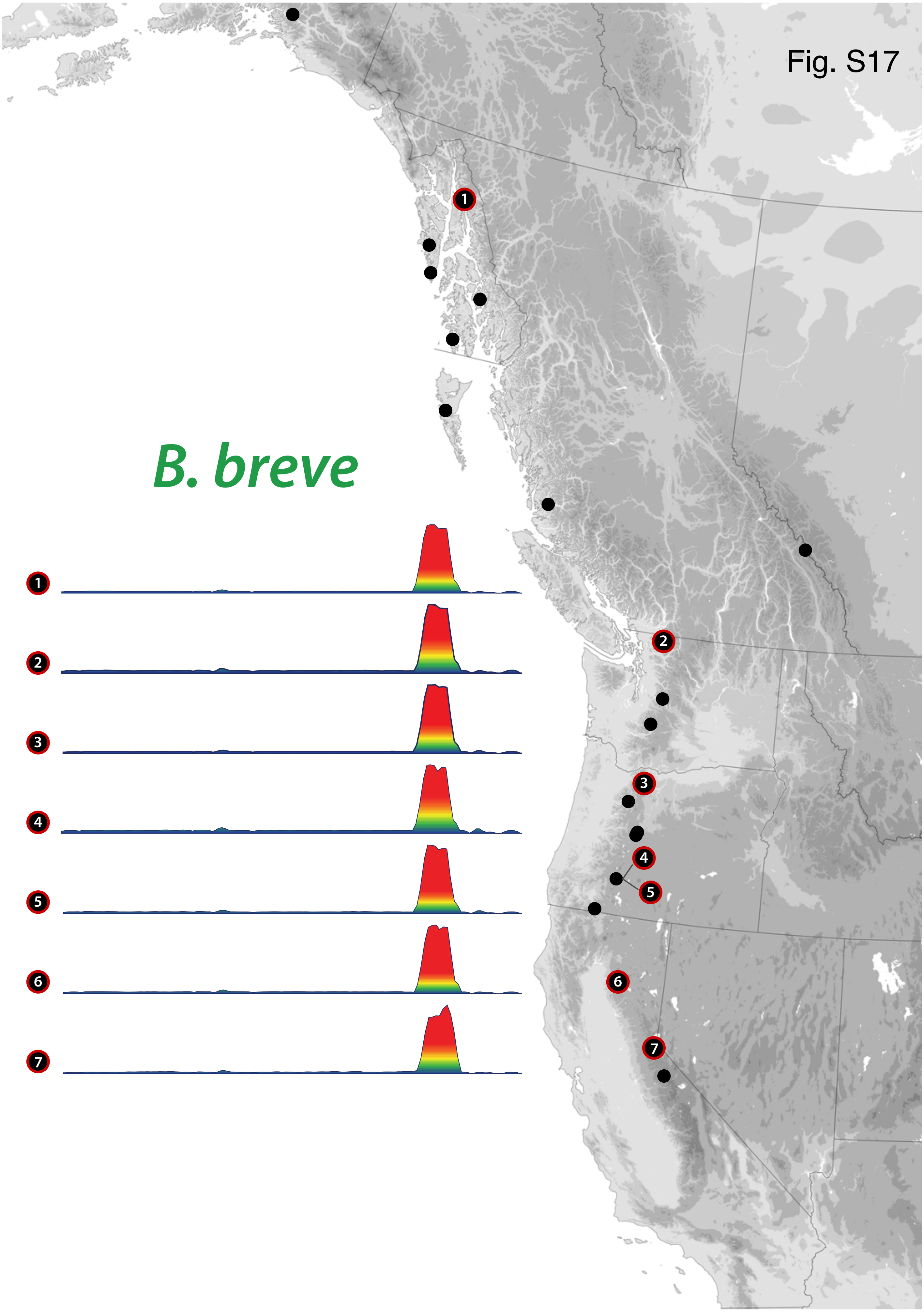

Fig. S17
B. breve
1
2
3
4
5
6
7

Fig. S18

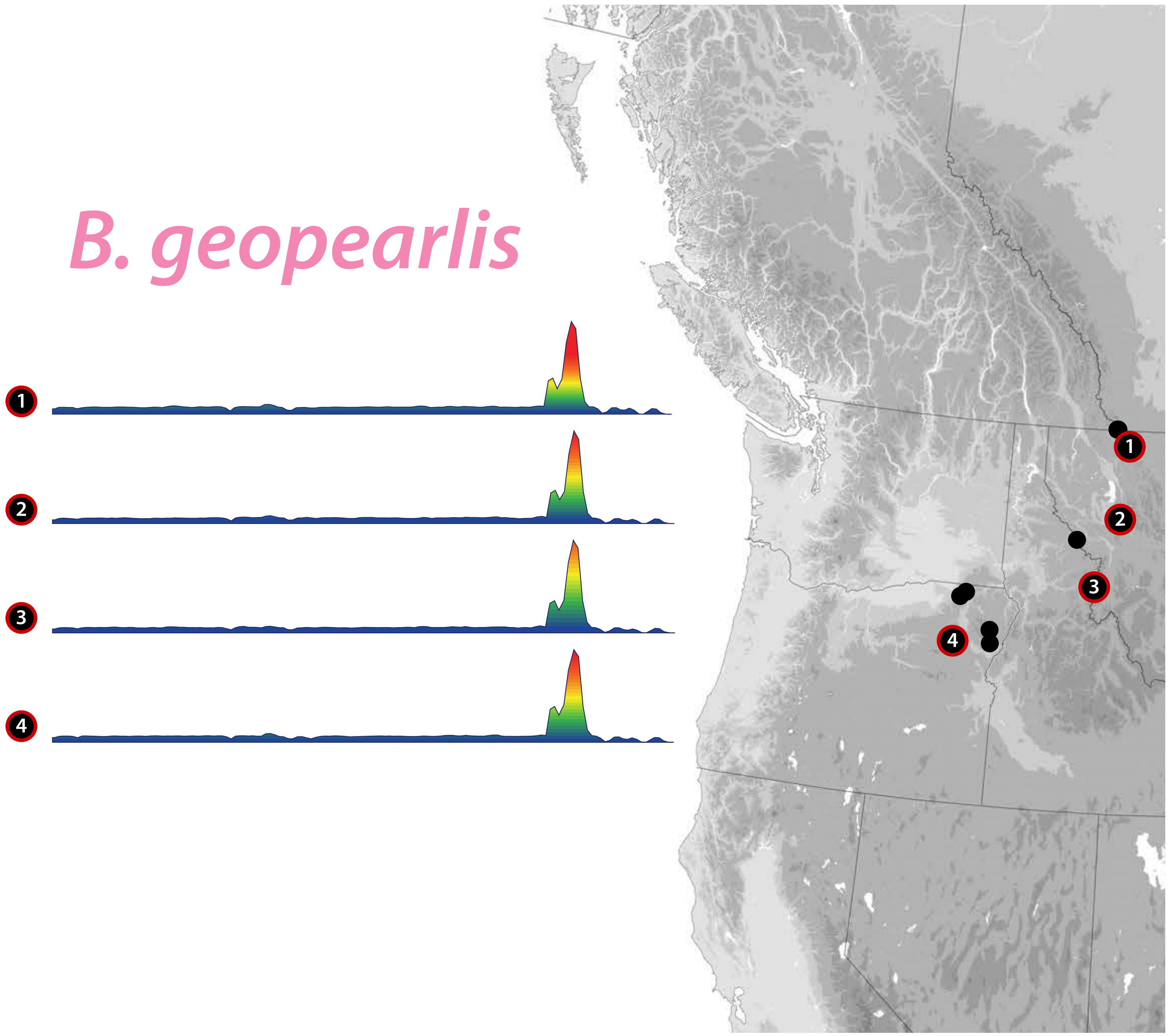

Fig. S19

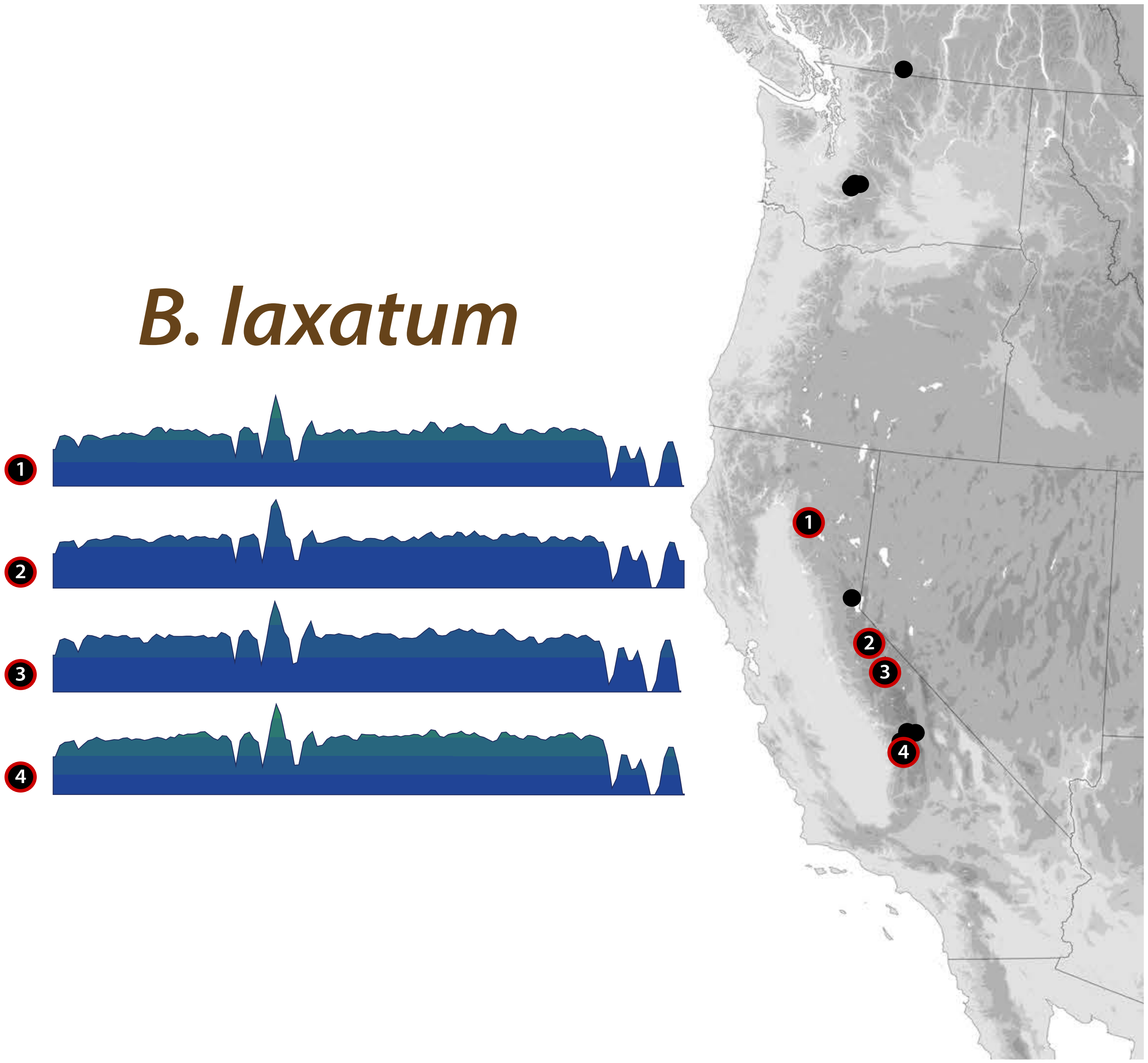

Fig. S20

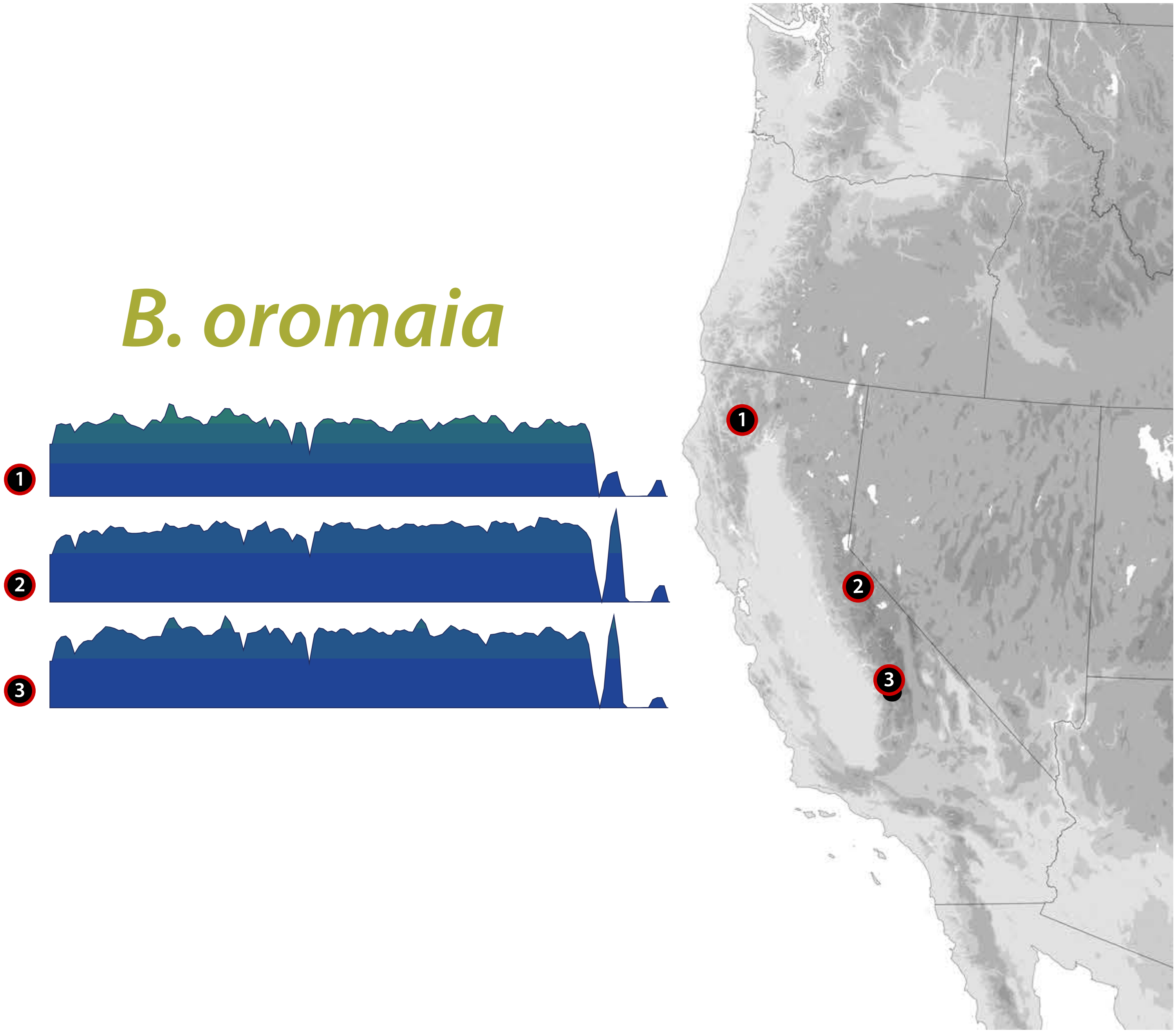

Fig. S21

#### *B. testatum*

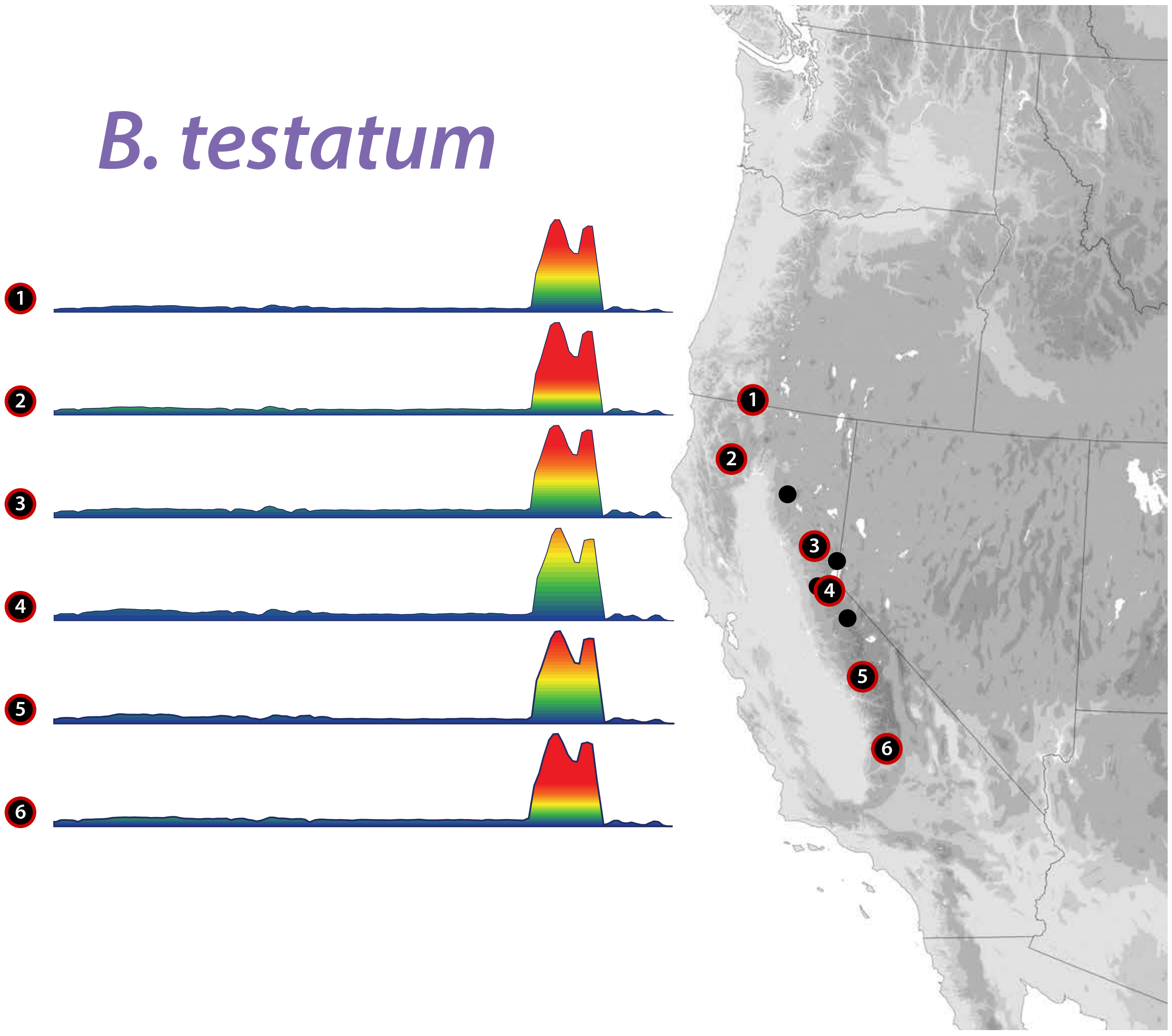

Fig. S22

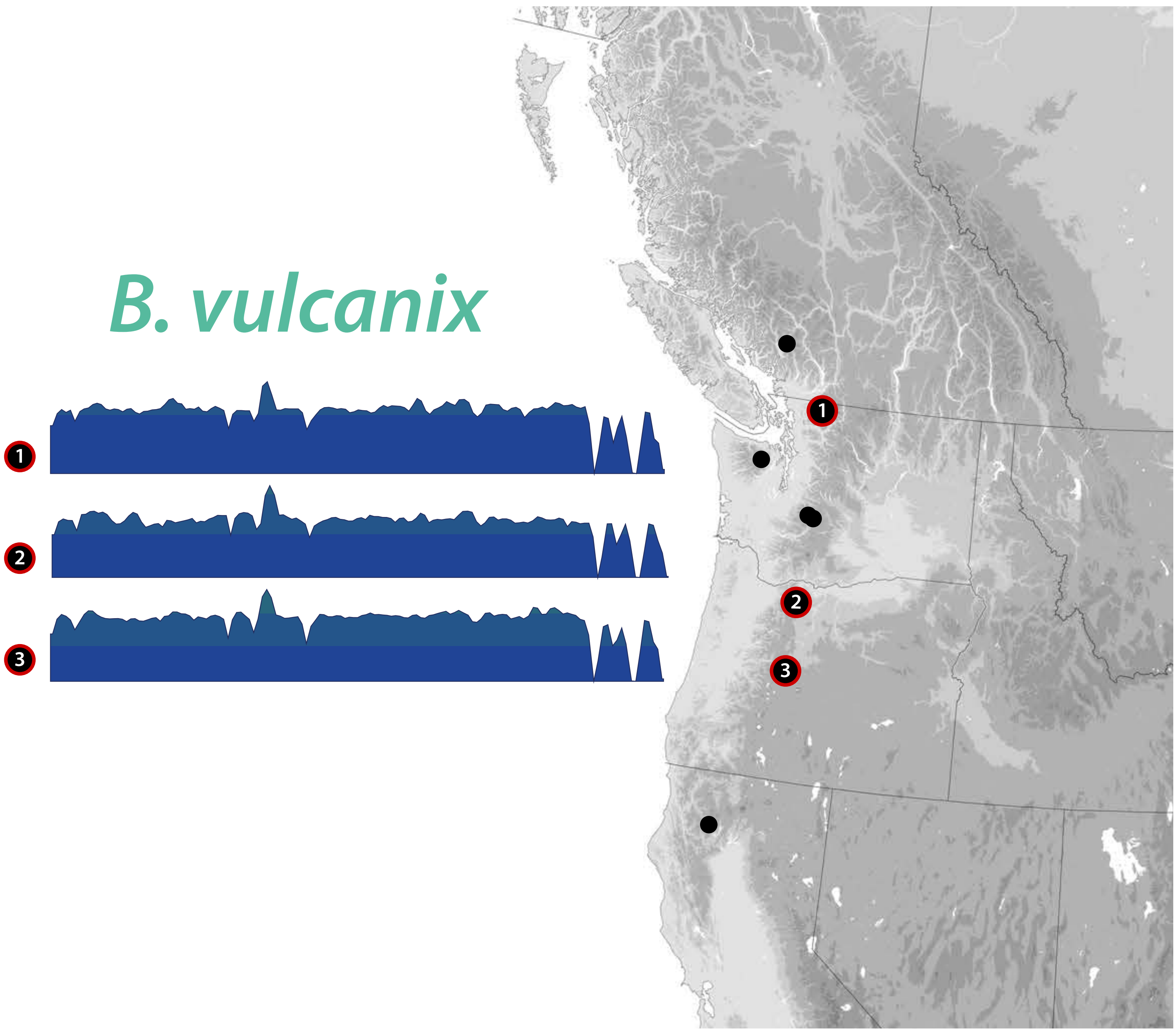

Fig. S23

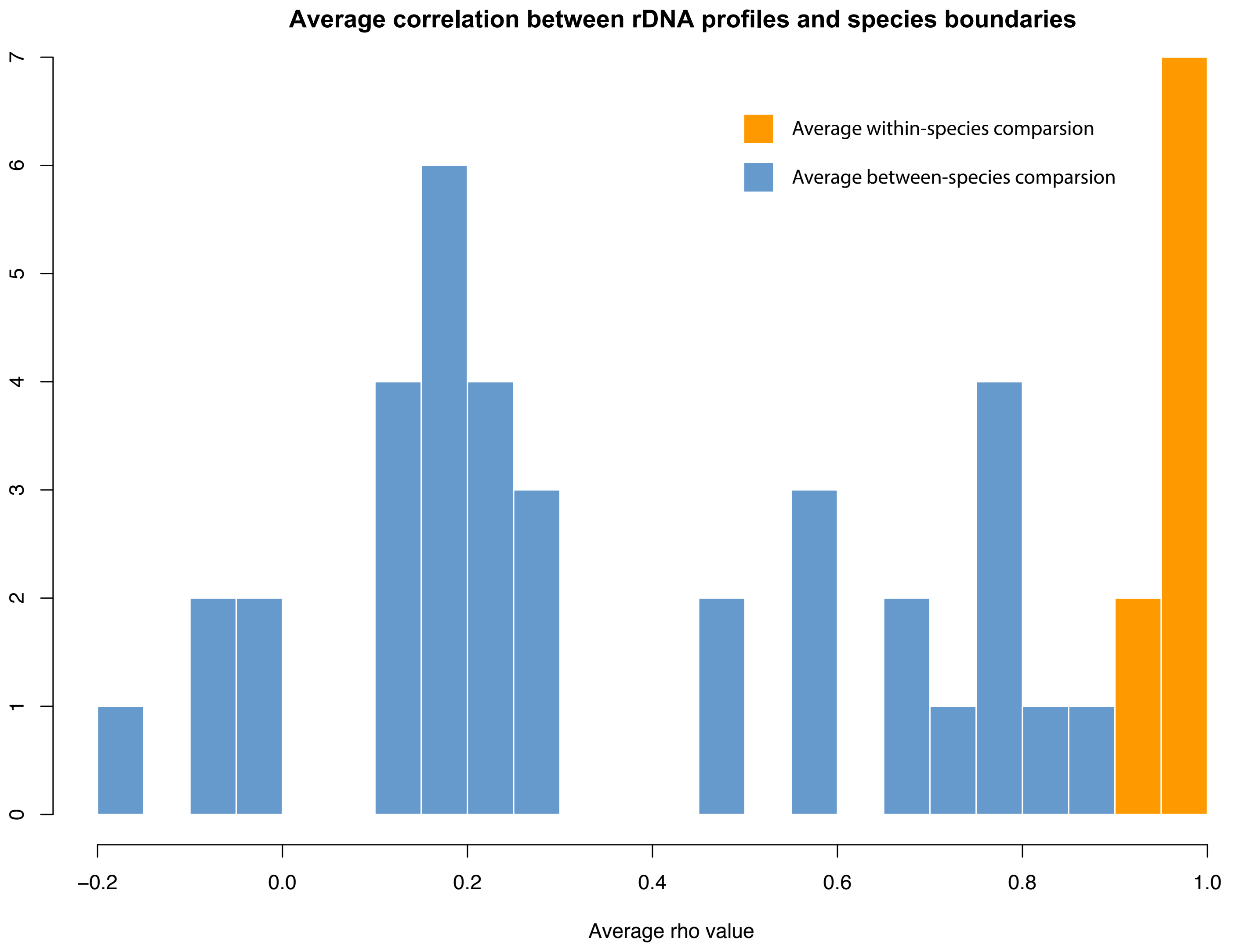

Average correlation between rDNA profiles and species boundaries
Average within-species comparsion
Average between-species comparsion
0
1
2
3
4
5
6
7
−0.2
0.0
0.2
0.4
0.6
0.8
1.0
Average rho value

Fig. S24

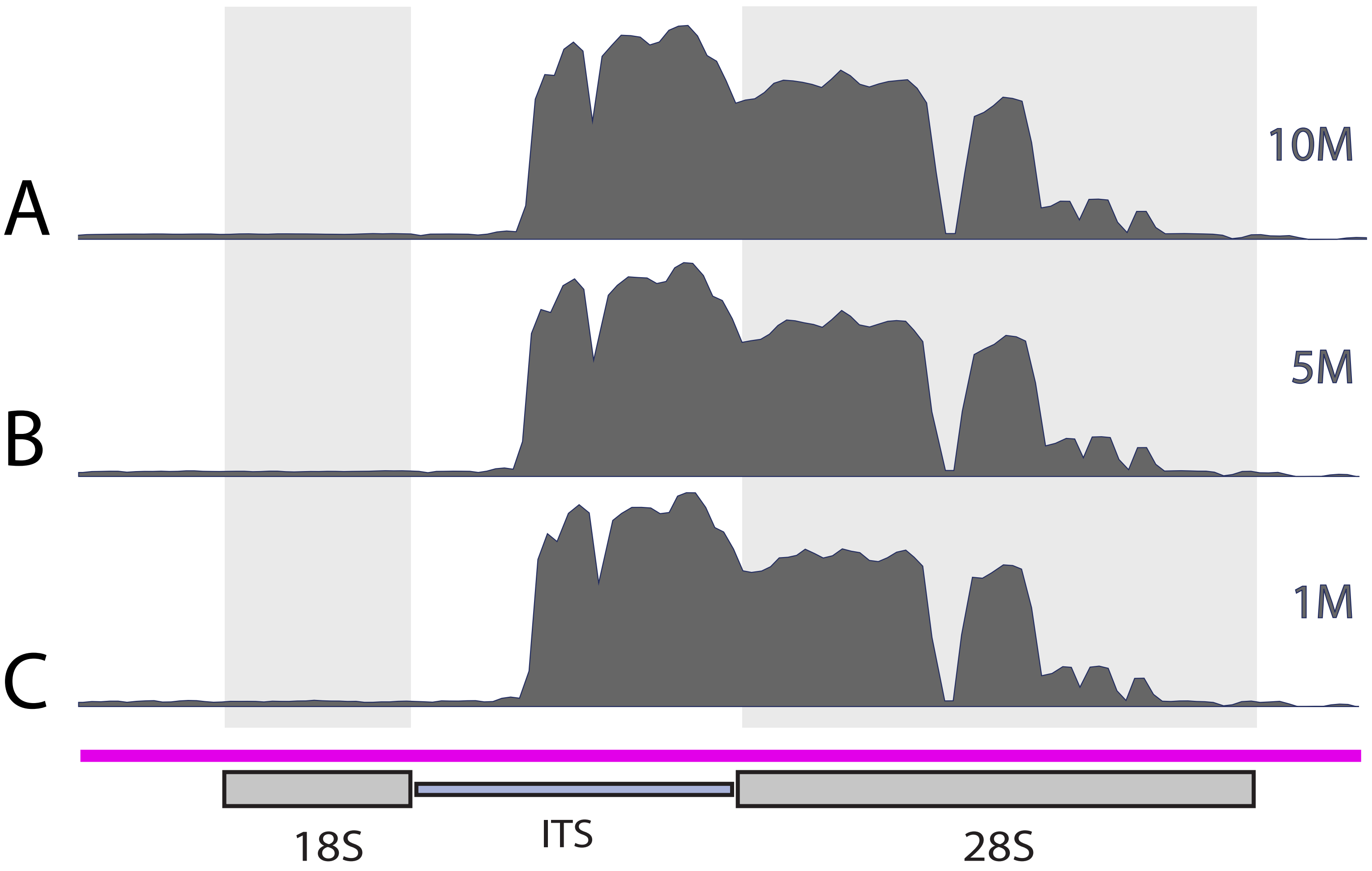

Fig. S25

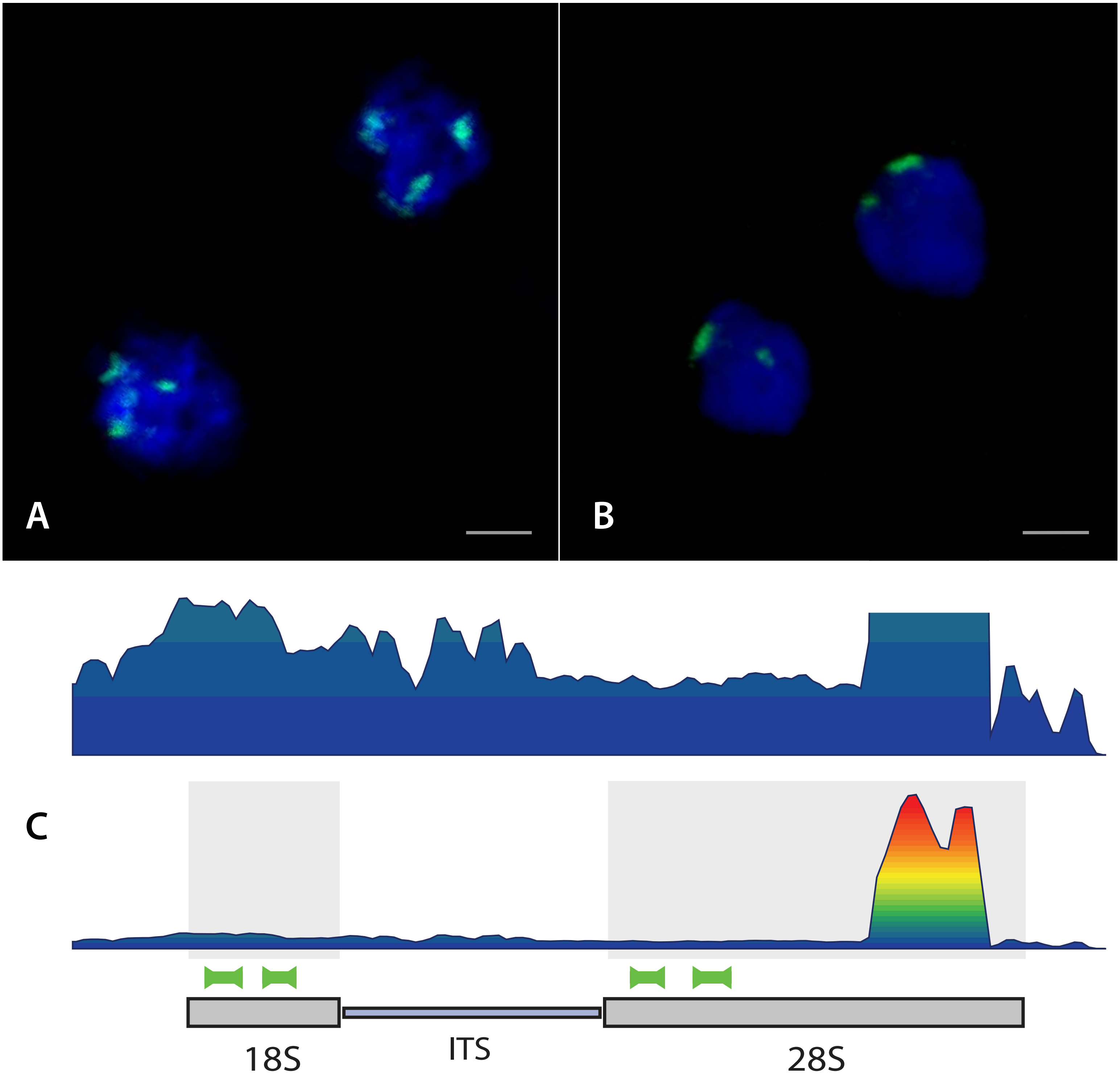

Fig. S26

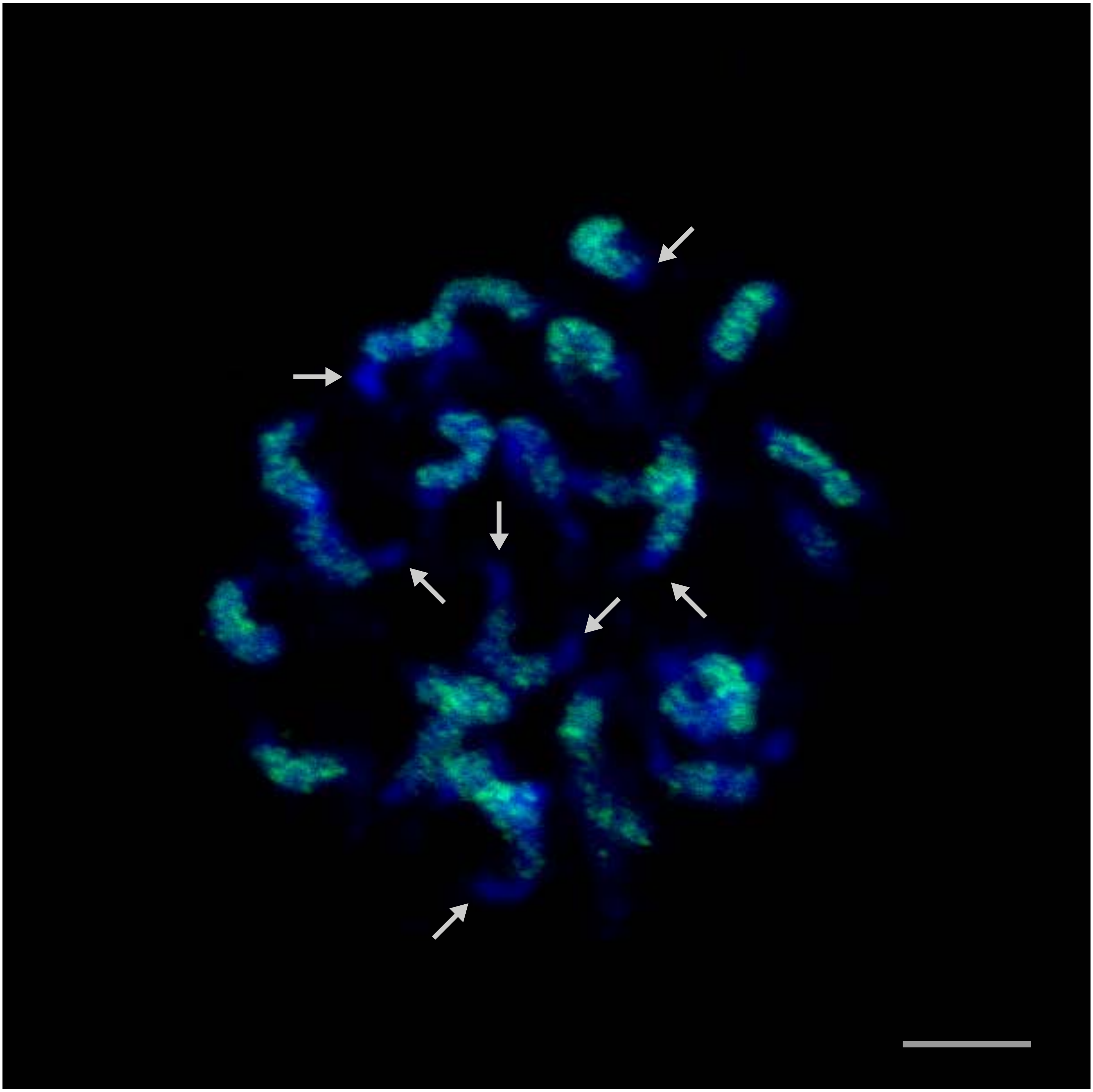

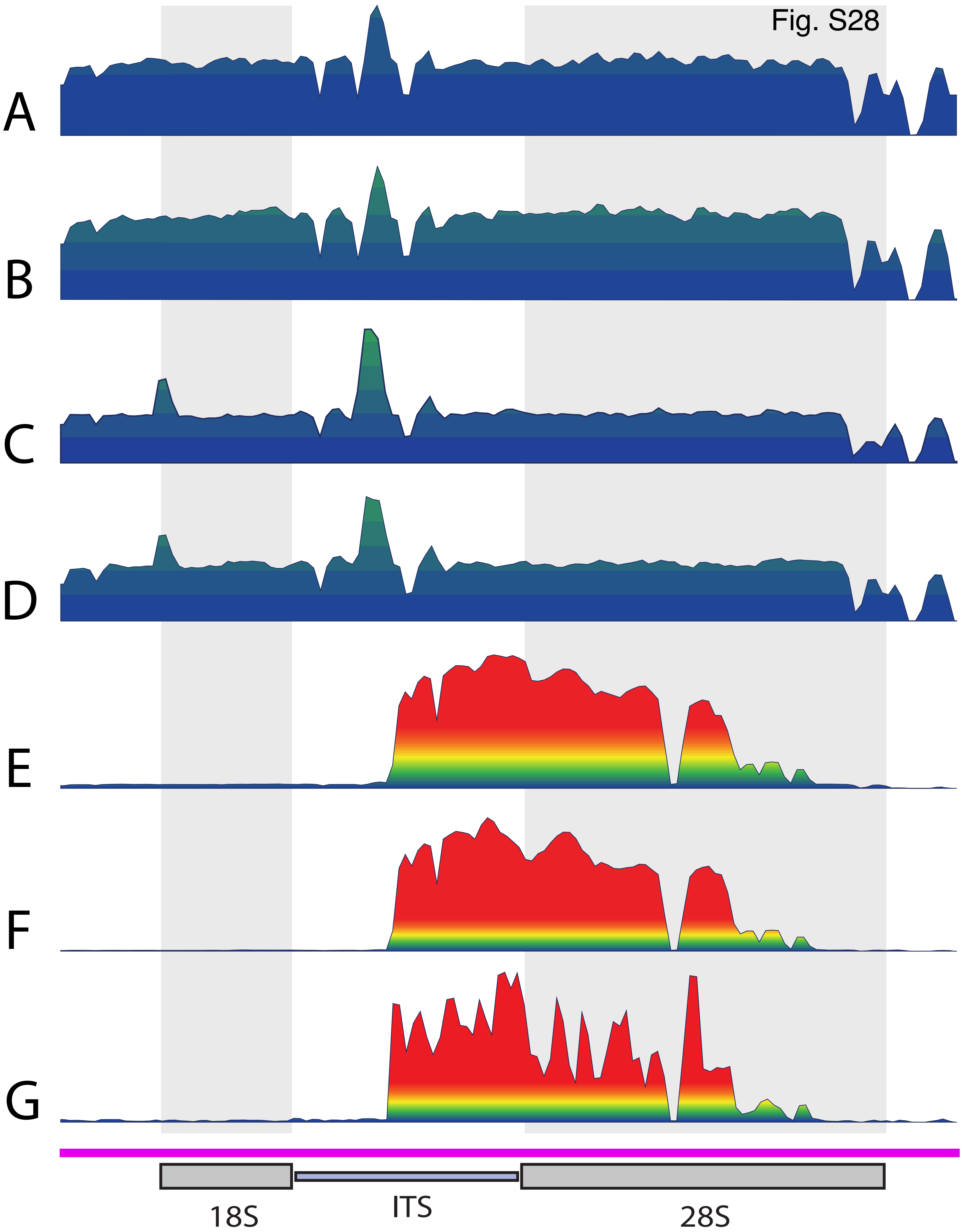

Fig. S28
A
B
C
D
E
F
G
18S
ITS
28S
